# ACE2-independent sarbecovirus cell entry is supported by TMPRSS2-related enzymes and reduces sensitivity to antibody-mediated neutralization

**DOI:** 10.1101/2024.04.18.590061

**Authors:** Lu Zhang, Hsiu-Hsin Cheng, Nadine Krüger, Bojan Hörnich, Luise Graichen, Alexander S. Hahn, Sebastian R. Schulz, Hans-Martin Jäck, Metodi V. Stankov, Georg M.N. Behrens, Marcel A. Müller, Christian Drosten, Onnen Mörer, Martin Sebastian Winkler, ZhaoHui Qian, Stefan Pöhlmann, Markus Hoffmann

## Abstract

The COVID-19 pandemic, caused by SARS-CoV-2, demonstrated that zoonotic transmission of animal sarbecoviruses threatens human health but the determinants of transmission are incompletely understood. Here, we show that most spike (S) proteins of horseshoe bat and Malayan pangolin sarbecoviruses employ ACE2 for entry, with human and raccoon dog ACE2 exhibiting broad receptor activity. The insertion of a multibasic cleavage site into the S proteins increased entry into human lung cells driven by most S proteins tested, suggesting that acquisition of a multibasic cleavage site might increase infectivity of diverse animal sarbecoviruses for the human respiratory tract. In contrast, two bat sarbecovirus S proteins drove cell entry in an ACE2-independent, trypsin-dependent fashion and several ACE2-dependent S proteins could switch to the ACE2-independent entry pathway when exposed to trypsin. Several TMPRSS2-related cellular proteases but not the insertion of a multibasic cleavage site into the S protein allowed for ACE2-independent entry in the absence of trypsin and may support viral spread in the respiratory tract. Finally, the pan-sarbecovirus antibody S2H97 enhanced cell entry driven by two S proteins and this effect was reversed by trypsin. Similarly, plasma from quadruple vaccinated individuals neutralized entry driven by all S proteins studied, and use of the ACE2-independent, trypsin-dependent pathway reduced neutralization sensitivity. In sum, our study reports a pathway for entry into human cells that is ACE2-independent, supported by TMPRSS2-related proteases and associated with antibody evasion.

## Introduction

The zoonotic transmission of animal coronaviruses of the genus *Betacoronavirus* to humans can present a major health threat. Thus, the transmission of SARS-CoV-1 from bats to humans via raccoon dogs and other intermediate hosts in 2002 resulted in the SARS epidemic that claimed roughly 800 lives ^1–3^. In 2012, a new, severe respiratory disease, Middle East respiratory syndrome (MERS), emerged in Saudi Arabia and was found to be caused by a novel coronavirus, MERS-CoV, which is transmitted from dromedary camels to humans and causes fatal diseases in roughly 30% of the afflicted patients ^4,5^. Finally, the emergence of SARS-CoV-2 in the human population in the winter season of 2019 in Hubei province, China, resulted in the COVID-19 pandemic that has claimed 18 million lives in the first two years alone ^6–8^. Although emergence of SARS-CoV-2 from a research laboratory has been suggested, a constantly increasing amount of evidence indicates that the virus was transmitted from animals to humans, likely at the Huanan Seafood market, in Wuhan, China ^9–11^. Thus, several betacoronaviruses from animals have zoonotic and pandemic potential and identifying which determinants control their ability to infect human cells will be instrumental for risk assessment and for devising antiviral strategies.

Trimers of the coronavirus spike protein (S) are incorporated into the viral envelope and facilitate viral entry into target cells. For this, the surface unit, S1, of S protein monomers binds to cellular receptors, ACE2, in case of SARS-CoV-1 and SARS-CoV-2 ^8,12,13^, while the S2 subunit facilitates fusion of the viral and a target cell membrane, allowing delivery of the viral genetic information into the host cell cytoplasm, the site of coronavirus replication. Cleavage of the S protein by host cell proteases at the S1/S2 site (located at the S1/S2 interface) and the S2’ site (located within the S2 subunit) is essential for membrane fusion and can be facilitated by the lysosomal cysteine protease cathepsin L or the cell surface serine protease TMPRSS2 ^12,14^ with the latter being essential for lung cell infection and pathogenesis ^14–17^. Finally, protease and receptor usage are major determinants of coronavirus cell and species tropism and are thus in the focus of many current research efforts ^18^.

The subgenus *Sarbecovirus* within the genus *Betacoronavirus* contains a single species, severe acute respiratory syndrome-related coronavirus. This species comprises SARS-CoV-2 and more than 100 related viruses that have been identified in bats and pangolins. A subset of these viruses can use angiotensin-converting enzyme 2 (ACE2) for entry into human and animal cells ^19–29^. However, it is not fully clear whether certain animal species can be identified as potential reservoirs or intermediate hosts for animal sarbecoviruses based on exceptionally broad receptor activity of their ACE2 orthologues. Recent studies provided evidence that the exposure of certain sarbecovirus S proteins to trypsin can facilitate ACE2-independent viral entry into human cells, a process that is determined by the receptor binding domain (RBD), and that equipping these S proteins with a multibasic cleavage site, a major virulence determinant of SARS-CoV-2 ^16,30^, is insufficient for trypsin-independent entry ^21,31–33^. However, these analyses were confined to small numbers of S proteins and it has not been resolved whether cellular proteases other than trypsin can cleave and activate trypsin-dependent sarbecovirus S proteins for host cell entry. Finally, it is incompletely understood whether antibodies elicited by multiple COVID-19 vaccinations neutralize a broad spectrum of animal sarbecoviruses and it is unknown how usage of the ACE2-independent pathway impacts susceptibility to antibody-mediated neutralization.

Here, examining a panel of bat and pangolin sarbecovirus S proteins, we found that multiple S proteins utilized human ACE2 for entry and that, among animal ACE2 orthologues, raccoon dog ACE2 exhibited the broadest receptor activity. We confirm that certain S proteins mediate ACE2-independent, trypsin-dependent entry and that this process is controlled by the RBD. Furthermore, we found that expression of certain type II transmembrane serine proteases (TTSPs) in particle-producing cells, analogous to trypsin treatment, allowed for ACE2-independent entry into human cells. In addition, we discovered that antibodies from quadruple vaccinated individuals neutralized entry driven by all S proteins studied, suggesting that COVID-19 vaccines might also offer some protection against diverse animal sarbecoviruses. Finally, we obtained evidence that ACE2-independent, trypsin-dependent entry can modulate neutralization by the pan sarbecovirus antibody S2H97 and allows for partial antibody evasion in the context of plasma from COVID-19 vaccinees.

## Results

### The RBMs of clade 2 and 4 sarbecovirus S proteins display major structural differences compared to clade 1 and 3 RBMs due to sequence variations in two surface exposed loops

The alignment of the amino acid sequence of 184 sarbecovirus S proteins revealed clustering into 5 clades and 14 S proteins, representing all clades, were selected for detailed analyses (Figure 1A and Supplemental figure 1). Structural studies had previously determined that the SARS-CoV-1 S (SARS-1-S) and SARS-CoV-2 S (SARS-2-S) receptor binding motifs (RBM), which are located within the RBD and make direct contact with ACE2, exhibit a similar structure ^34^. The predicted structures of the RBMs of bat sarbecovirus clade 1 S proteins were similar among each other and comparable to that of the RBM of SARS-1-S, the prototypic clade 1 S protein (Figure 1B and Supplemental figure 2). Similar findings were made for the structures of clade 3 RBMs, including the RBM of the SARS-2-S (Figure 1B). In contrast, loop 1 in the RBM was largely absent from clade 2 and clade 4 S proteins (Figure 1B-C and Supplemental figure 2) and some clade 4 S proteins contained a shortened loop 2 (Figure 1B-C and Supplemental figure 2). Thus, the RBMs of the S proteins selected for analysis likely exhibit similar structures but two surface exposed loops are partially or largely absent from clade 2 and 4 S proteins, due to clade-specific sequence variations in the S gene, which may impact receptor interactions.

**Fig. 1.**
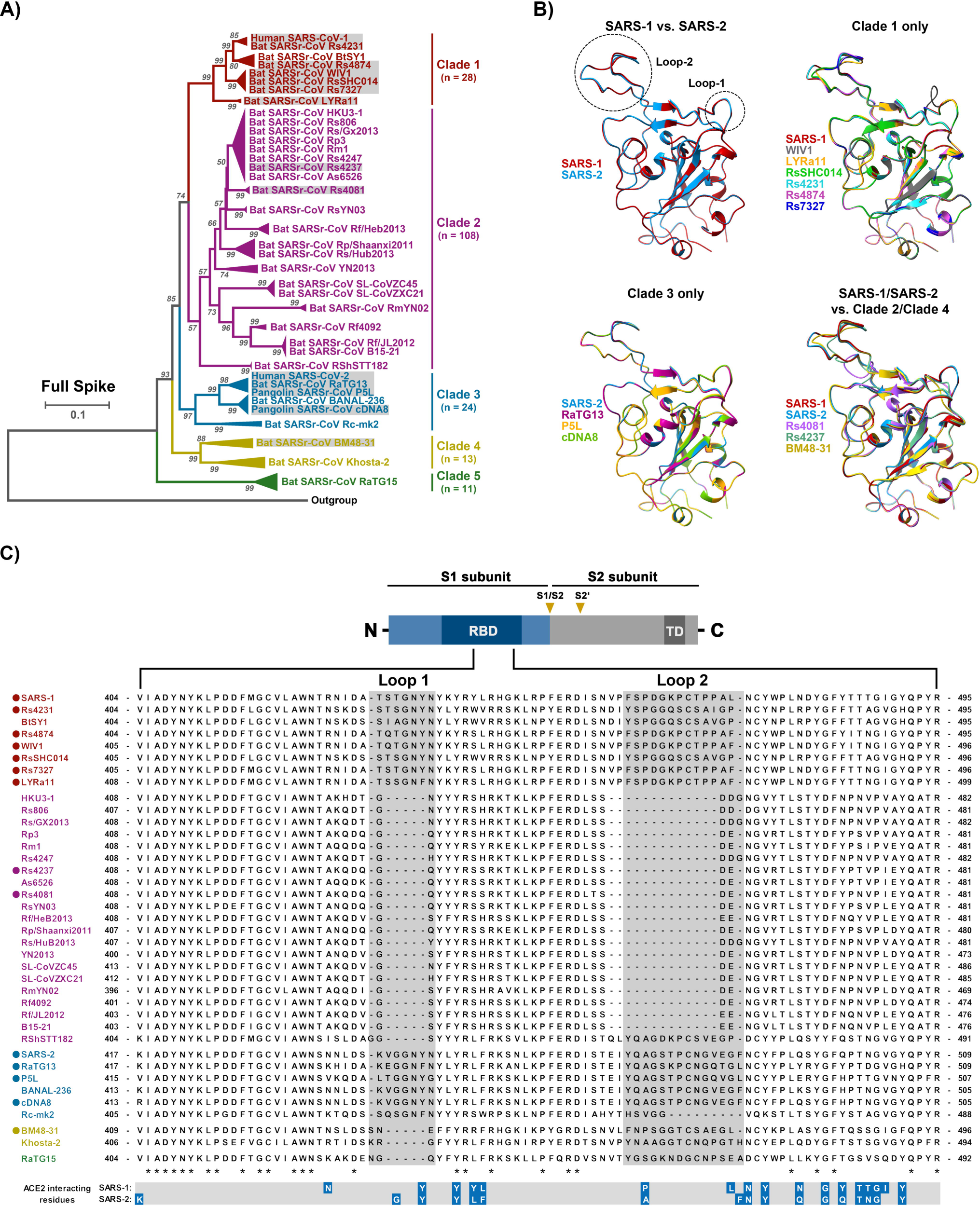
Alignment of S protein sequences and structural predictions. **A**) Phylogenetic analysis of human and animal sarbecoviruses. The sarbecoviruses were grouped into five clades, indicated by different colors, based on the full spike sequences. The sarbecoviruses functionally analyzed in the present study are indicated in grey boxes. (See Figure S1 for more details). **B**) Structure of RBD. The structure of RBDs was predicted based on homology modeling using SARS-2-S RBD as template. Two loops involved in ACE2 interactions are highlighted (See supplemental figure 2 for more details). **C**) Schematic overview of the spike (S) protein domain structure (upper panel) and alignment of the RBM sequences of the S proteins analyzed in panel A. The ACE2 interacting residues of SARS-1-S and SARS-2-S are marked in blue (lower panel). “∗” indicates conserved amino acid residues, “-“ indicates gaps. The S proteins under study are indicated by circles. Abbreviations: NTD = N-terminal domain; RBD = receptor-binding domain; TD = transmembrane domain; S1/S2 and S2’ = cleavage sites in the S protein.

### Raccoon dog ACE2 exhibits broad receptor activity for clade 1 and 3 animal sarbecoviruses

For a detailed analysis of determinants governing entry of animal sarbecoviruses into human cells we selected a total of 14 S proteins that represent sarbecovirus clades 1 to 4, including SARS-1-S (clade 1, human), WIV1 (clade 1, bat), LYRa11 (clade 1, bat), RsSHC014-S (clade 1, bat), Rs4231 (clade 1, bat), Rs4874 (clade 1, bat), Rs7327 (clade 1, bat), Rs4081-S (clade 2, bat), Rs4237 (clade, bat), SARS-2-S (clade 3, human), RaTG13-S (clade 3, bat), P5L-S (clade 3, Malayan pangolin), cDNA8-S (clade 3, Malayan pangolin), and BM48-31-S (clade 4, bat).

We first asked whether the sarbecovirus S proteins under study can employ human ACE2 and ACE2 orthologues from animal species potentially relevant to zoonotic transmission for cell entry. For this, we studied receptor activity of human, pig, mink, pangolin and bat ACE2, using pseudotyped particles and transfected BHK-21 hamster cells. BHK-21 cells were chosen as targets since they do not support ACE2-dependent entry due lack of ACE2 expression ^24^. All ACE2 orthologues analyzed were robustly expressed in transfected cells, as determined by immunoblot (data not shown). The analysis of binding of soluble human ACE2 to S protein expressing cells revealed robust ACE2 binding to cells expressing SARS-2-S, SARS-1-S, WIV1, LYRa11 S protein while for several other S proteins, including RaTG13 S protein, moderate to low ACE2 binding was observed (Figure 2A and Supplemental figure 3). Finally, cells expressing Rs4081, Rs4237 and BM48-31 S proteins failed to bind to soluble ACE2 (Figure 2A).

**Fig. 2.**
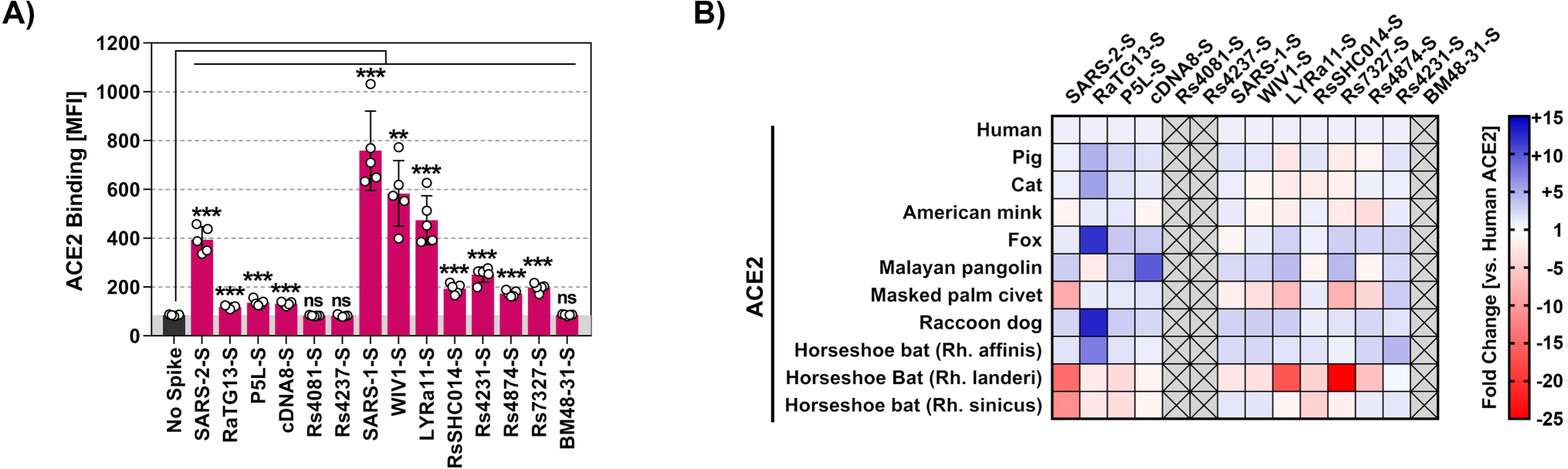
Raccoon dog ACE2 supports entry driven by the S proteins of diverse sarbecoviruses. **A**) Binding of soluble human ACE2 to S protein expressing cells. 293T cells transiently expressing the indicated S proteins (or no S protein) were first incubated with soluble ACE2 containing a C-terminal Fc-tag (derived from human immunoglobulin G; solACE2-Fc) and subsequently incubated with an AlexaFluor-488-coupled secondary antibody, before solACE2-Fc binding was analyzed by flow cytometry (see Figure S3 for details on the gating strategy). Presented are the average (mean) data from five biological replicates (each conducted with single samples) in which solACE2-Fc binding to S protein expressing cells was normalized against binding to control transfected cells (set as 100). Error bars indicate SEM. Statistical significance was assessed by two-tailed Student’s t-tests (p > 0.05, not significant [ns]; p ≤ 0.05, *; p ≤ 0.01, **; p ≤ 0.001, ***). **B**) Receptor activity of ACE2 orthologues. BHK-21 cells transiently expressing the indicated ACE2 orthologues (or empty vector) were inoculated with pseudotyped particles bearing the indicated S proteins (or no S protein). Entry into cells expressing ACE2 orthologues was normalized against entry into cells expressing human ACE2 (set as 1). The heat map presents the average (mean) data from three biological replicates (each conducted with four technical replicates). (See Figure S4 for more details)

Next, we determined whether the S proteins could employ ACE2 of human and animal origin for host cell entry. The S proteins that bound to human ACE2 were able to use human ACE2 and animal ACE2 orthologues for entry into transfected BHK-21 cells but differences in breadth of receptor activity were noted (Figure 2B and supplemental figure 4). Thus, all S proteins studied efficiently employed human ACE2 for entry with the exception of the aforementioned S proteins of BM48-31, Rs4081 and Rs4237, which had also failed to bind to ACE2 (Figure 2B). Further, ACE2 usage by cDNA8 S protein was generally inefficient and ACE2 of the Lander’s horseshoe bat, (*Rhinolophus landeri*), did not appreciably support entry driven by the S proteins of LYRa11, RsSHC014 and Rs7327 although these S proteins could use other ACE2 orthologues for robust entry (Figure 2B). Finally, a systematic comparison of all ACE2 orthologues revealed that ACE2 from the raccoon dog supported entry driven by all tested clade 1 and 3 sarbecovirus S proteins with at least the same or, for several S proteins, even higher efficiency than human ACE2 (Figure 2B), in keeping with a potential role of raccoon dogs as intermediate host or reservoir for several animal sarbecoviruses.

We next asked whether the S proteins analyzed were able to mediate entry into diverse human and animal cell lines. For this, we used 293T cells (human, kidney), 293T cells engineered to overexpress human ACE2, Vero cells (African green monkey, kidney), Vero cells engineered to overexpress TMPRSS2 or TMPRSS2 jointly with ACE2, A549 cells (human, lung) engineered to overexpress human ACE2 and TMPRSS2, Calu-3 (human, lung), Calu-3 cells engineered to overexpress human ACE2, Caco-2 cells (human, colon) and Huh-7 cells (liver, human) as targets.

All cell lines expressed endogenous or exogenous ACE2 and thus allowed for SARS-CoV-2 S protein-driven entry (Figure 3A). Most animal sarbecovirus S proteins mediated entry into cell lines expressing endogenous ACE2 (Figure 3A) and entry was markedly increased upon directed expression of ACE2 in 293T and Calu-3 cells (Figure 3A). In contrast, directed expression of TMPRSS2 in Vero cells had only moderate effects on viral entry. Thus, most S proteins tested were able to bind to human ACE2, although with different efficiencies, and were able to mediate entry into cell lines expressing human ACE2 or animal ACE2 orthologues. In contrast, three S proteins, BM48-31, Rs4237 and Rs4081, failed to mediate entry into any of the cell lines tested, irrespective of ACE2 expression (Figure 3A) and were thus in the focus of our further analyses.

**Fig. 3.**
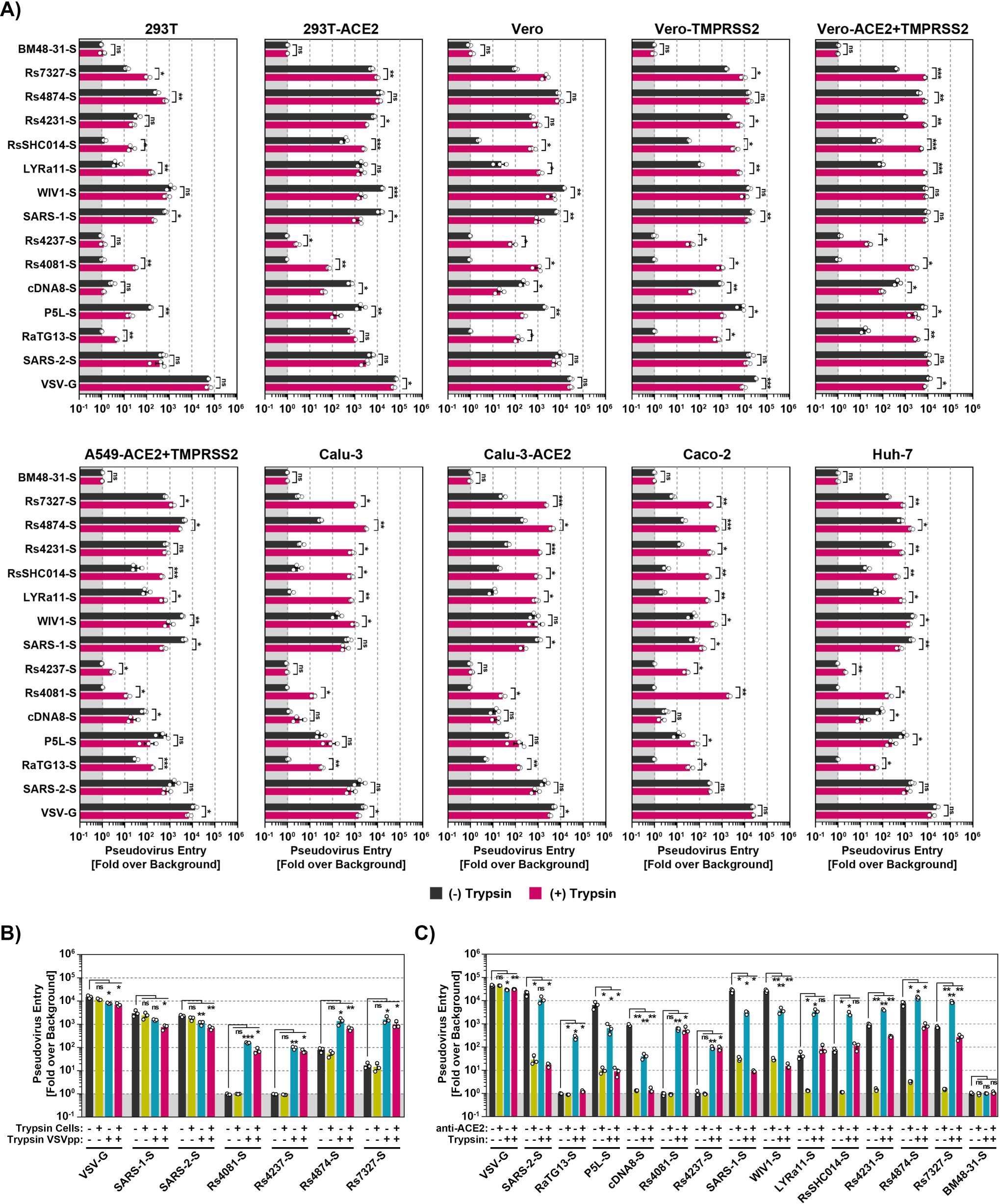
Trypsin treatment can allow for ACE2-independent cell entry. **A**) S protein driven cell entry in the presence and absence of trypsin. Particles bearing the indicated S proteins (or no S protein) were preincubated with or without trypsin before being added to the respective cell lines. S-protein driven cell entry was analyzed by measuring the activity of virus-encoded firefly luciferase in the cell lysate at 16-18h post inoculation. Presented are the average of (mean) data from three biological replicates (each conducted with four technical replicates) in which cell entry was normalized against that measured for particles bearing no S protein (set as 1). Error bars show the SEM. Statistical significance was assessed by two-tailed Student’s t-tests (p > 0.05, not significant [ns]; p ≤ 0.05, *; p ≤ 0.01, **; p ≤ 0.001, ***). **B**) Trypsin treatment of viral particles but not target cells promotes entry. Vero cells or pseudotyped particles bearing indicated S proteins were pre-incubated with trypsin and subsequently trypsin inhibitor as indicated. The pseudotyped particles were added to the cells. S-protein-driven cell entry was analyzed by and data presented as described for panel A. Presented are the average (mean) data of three biological replicates, each performed with four technical replicates. Error bars show SEM. Statistical significance was assessed by ANOVA (p > 0.05, not significant [ns]; p ≤ 0.05, *; p ≤ 0.01, **; p ≤ 0.001, ***). **C**) Blockade of S protein-driven cell entry by an anti-ACE2 antibody. Vero-TMPRSS2 cells were pre-incubated with anti-ACE2 antibody. Particles bearing the indicated S proteins were incubated with trypsin followed by incubation with trypsin inhibitor before addition onto target cells. The pseudotyped particles were added to the cells. S-protein-driven cell entry was analyzed by and data presented as described for panel A. Presented are the average (mean) data of three biological replicates, each performed with four technical replicates. Error bars show SEM. Statistical significance was assessed by ANOVA (p > 0.05, not significant [ns]; p ≤ 0.05, *; p ≤ 0.01, **; p ≤ 0.001, ***).

### The S proteins of clade 2 bat sarbecoviruses Rs4237 and Rs4081 mediate trypsin-dependent entry into human cells

We next asked whether lack of proteolytic activation of the BM48-31, Rs4237 and Rs4081 S proteins was responsible for lack of cell entry. To address this possibility, we preincubated S protein-bearing particles with trypsin before addition to target cells. Trypsin treatment modulated S protein-driven entry in a cell line- and S protein-dependent manner. For one group of S proteins, including the S proteins of Rs7327, Rs4231, RsSHC014, trypsin treatment either increased entry efficiency or had no impact (Figure 3A). For instance, entry of Rs4231 S protein into Calu-3 and Caco-2 cells was markedly increased by trypsin pre-treatment although this effect was not observed with 293T cells. For a second group of S proteins, including those of SARS-CoV-1, WIV1 and cDNA8, trypsin treatment reduced entry efficiency or did not change entry efficiency (Figure 3A). Interestingly, among the S proteins that were unable to mediate cell entry in the absence of trypsin, trypsin pre-treatment allowed for Rs4081 S protein-driven entry into all cell lines studied (Figure 3A) with bat-derived MyDauLu/47 cells being the only exception (Supplemental figure 5). Similarly, trypsin pre-treatment allowed for Rs4237 S protein-driven entry into most cell lines studied, except for 293T, Calu-3 cell lines (Figure 3A) and most bat-derived cell lines studied (Supplemental figure 5). In contrast, BM48-31 S protein failed to mediate entry into any of the cell lines tested even upon pre-treatment with trypsin (Figure 3A and Supplemental figure 5). In sum, availability of an appropriate protease can limit sarbecovirus entry into human cells and this limitation can be overcome by trypsin treatment, in keeping with published data ^21,31,35,36^.

We next investigated whether trypsin promoted viral entry by acting on viral particles or on target cells. For this, cells, particles or particles and cells were preincubated with trypsin followed by addition of a trypsin inhibitor and mixing of particles and cells. Treatment of target cells with trypsin had no effect on entry driven by VSV-G or any of the sarbecovirus S proteins studied (Figure 3B). In contrast, pretreatment of particles with trypsin allowed for entry driven by the Rs4081 and Rs4237 S proteins and augmented entry driven by Rs4874 and Rs7327 but not SARS-CoV-1 and SARS-CoV-2 S proteins (Figure 3B). Finally, augmentation of viral entry by trypsin treatment of particles was not further increased when both particles and target cells were preincubated with trypsin (Figure 3B), indicating that trypsin acts on viral particles rather than target cells to promote entry driven by a subgroup of sarbecovirus S proteins.

### Trypsin-dependent cell entry driven by the S proteins of Rs4237 and Rs4081 is ACE2-independent

The finding that trypsin-promoted entry driven by the Rs4081and Rs4237 S proteins did not correlate with ACE2 expression suggested that these S proteins might mediate entry in an ACE2-independent manner. In order to investigate this possibility, we pre-treated particles with trypsin and/or target cells with anti-ACE2 antibody before infection. The anti-ACE2-antibody blocked entry driven by most S proteins studied, and for several S proteins, including SARS-CoV-2 S protein, trypsin treatment did not alter the efficiency of entry inhibition by the ACE2 antibody (Figure 3C). However, for other S proteins that facilitated ACE2-dependent entry, including LYRa11, RsSHC014, Rs4231, Rs4874 and Rs7327, trypsin treatment reduced the inhibitory effect of the anti-ACE2 antibody (Figure 3C). Finally, treatment of cells with anti-ACE2 antibody did not block trypsin-dependent cell entry driven by Rs4081and Rs4237 S proteins (Figure 3C). Collectively, these results demonstrate that Rs4081 and Rs4237 S protein engage a receptor other than ACE2 for host cell entry and that trypsin treatment can confer partial ACE2-independence to entry driven by other S proteins, including LYRa11, RsSHC014, Rs4231, Rs4874 and Rs7327.

### Trypsin cleaves sarbecovirus S proteins

We next investigated whether trypsin treatment resulted in S protein cleavage and how much trypsin was needed for S protein cleavage and S protein-driven entry. For analysis of cleavage, S protein bearing VSV particles were incubated with 0.5, 5 and 50 µg/ml of trypsin and then analyzed by immunoblot. All S proteins were largely uncleaved in the absence of trypsin, as documented by prominent signals for the uncleaved S0 protein, with exception of SARS-2-S, which was efficiently cleaved in the absence of trypsin due to the presence of a unique furin cleavage site (Figure 4A). The addition of trypsin led to the cleavage of all S proteins studied, as indicated by a reduction in signals for the S0 protein and an increase in signals corresponding to the S2 subunit (Figure 4A). For some of the S proteins additional signals were observed in the presence of 50 µg/ml trypsin, which likely corresponded to the S2’ fragment (produced upon cleavage of the S protein at the S2’ site) and cleavage products thereof (Figure 4A). Thus, all S proteins studied were cleaved by trypsin, although with different efficiencies, resulting in a concentration-dependent disappearance of S0 and appearance of the S2’ fragment and S2’ sub-fragments.

**Fig. 4.**
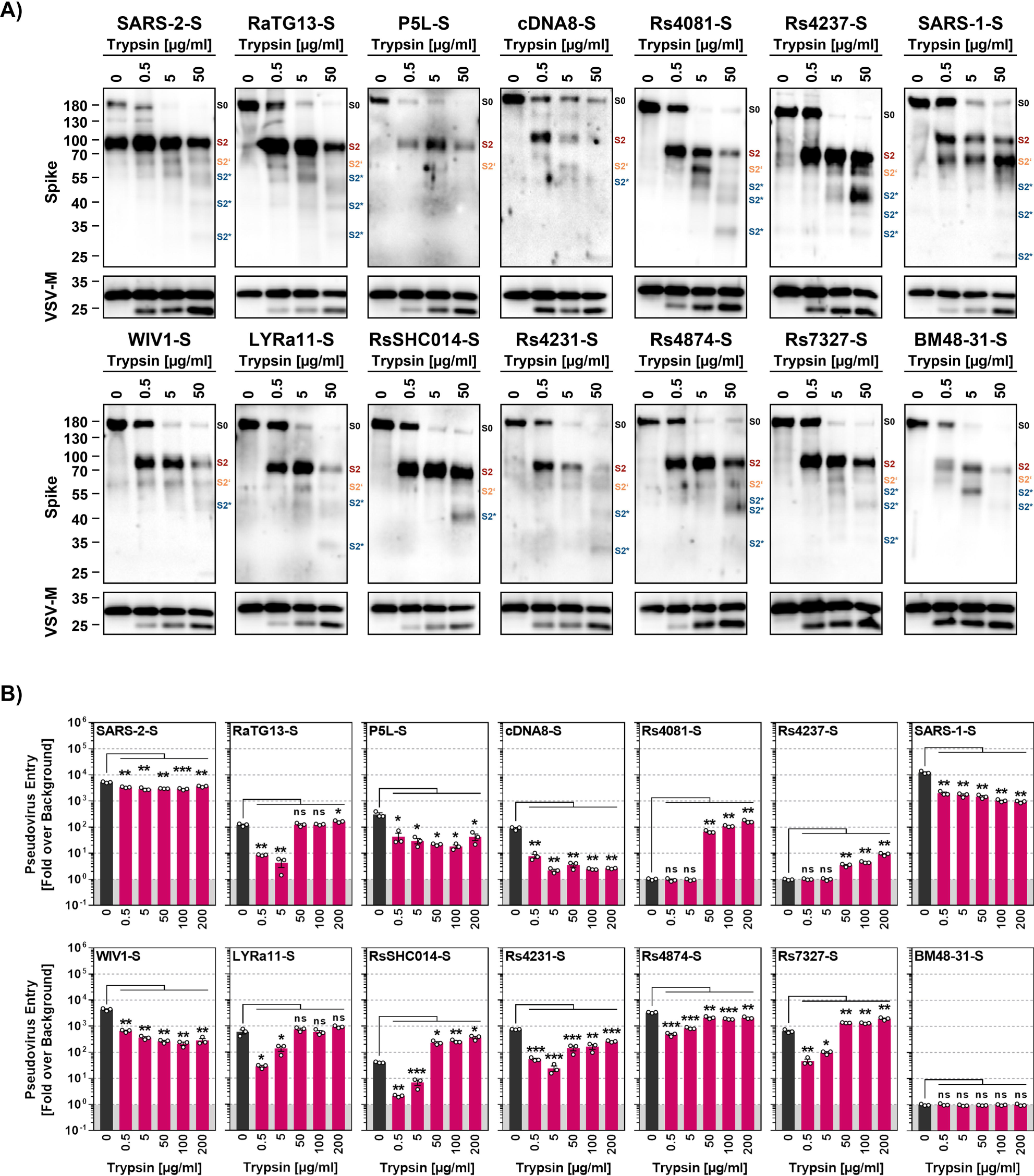
Trypsin cleaves the S proteins of diverse sarbecoviruses. **A**) Cleavage of S proteins by trypsin. Particles pseudotyped with the indicated S proteins were incubated with the indicated concentrations of trypsin for 20 min at 37℃ and S protein expression analyzed by immunoblot with SARS-CoV-2 S2 antibody. VSV-M served as loading control. Similar results were obtained in two separate experiments. Bands corresponding to uncleaved S proteins (S0), the S2 subunit (S2), S2 subunit cleaved at the S2’ site (S2’) and additional S2 cleavage fragments (S2*) are indicated. **B**) Modulation of S protein driven entry by trypsin is concentration-dependent. Particles pseudotyped with the indicated S proteins were treated with the indicated concentrations of trypsin for 20 min at 37L before addition to Vero cells. The efficiency of S protein-driven cell entry was determined by measuring the activity of virus-encoded firefly luciferase in cell lysates at 16-18h post inoculation. Results for S protein bearing particles were normalized against those obtained for particles bearing no S protein (set as 1). The average (mean) data of three biological replicates is presented, each performed with four technical replicates. Error bars show the SEM. Statistical significance was assessed by ANOVA (p > 0.05, not significant [ns]; p ≤ 0.05, *; p ≤ 0.01, **; p ≤ 0.001, ***).

We next analyzed concentration-dependence of trypsin-dependent S protein-driven cell entry by pre-incubation of pseudotyped particles with increasing amounts of trypsin. For this, we chose 200 µg/ml trypsin as maximal concentration, considering that concentrations of roughly 150 µg/ml are present in the human intestine ^37^. S proteins that did not exhibit augmented cell entry activity upon exposure to 50 µg/ml trypsin (Figure 3A), including SARS-1-S and SARS-2-S, were also not appreciably stimulated for augmented cell entry when a higher concentration of trypsin was used (Figure 4B). In contrast, S proteins that mediated increased entry upon exposure to 50 µg/ml trypsin, including RsSHC014 and RS7327 S proteins, were slightly more active in the presence of 200 µg/ml and this group included the S proteins of Rs4081 and Rs4237, which allowed for cell entry only upon trypsin-treatment (Figure 4B). Importantly, trypsin-treatment did not increase the ability of the S proteins to bind to ACE2 (Supplemental figure 6). Collectively, we found that 50 and 200 µg/ml trypsin robustly increased or allowed for cell entry activity of several animal sarbecovirus S proteins and these protease concentrations are likely attained in the intestine, which is believed to be a major target for sarbecovirus infection in bats ^38–40^. On a more general level, our findings suggest that lack of proteolytic activation of the viral S protein might impede host cell entry of Rs4237 and Rs4081.

### Thermolysin and elastase cleave Rs4081 S protein at the S1/S2 site and confer infectivity to Rs4081 S protein-bearing particles

We next investigated whether secreted proteases other than trypsin can promote entry driven by the Rs4081 S protein. For this, we first analyzed the effect of thermolysin, papain and elastase on cell entry. Thermolysin is a bacterial protease, while papain is a protease produced in plants, and thermolysin has been used previously to characterize coronavirus S proteins ^41^. Elastase promotes inflammation and plays a role in several lung pathologies, likely including COVID-19 ^42,43^. Immunoblot analyses revealed that trypsin, thermolysin and elastase cleaved both SARS-1-S and Rs4081-S at the S1/S2 site, resulting in production of the S2 fragment (Figure 5A). In contrast, papain digest of SARS-1-S and Rs4081-S resulted in several S2-derived fragments, suggesting multiple papain cleavage sites in the S2 subunit (Figure 5A).

**Fig. 5.**
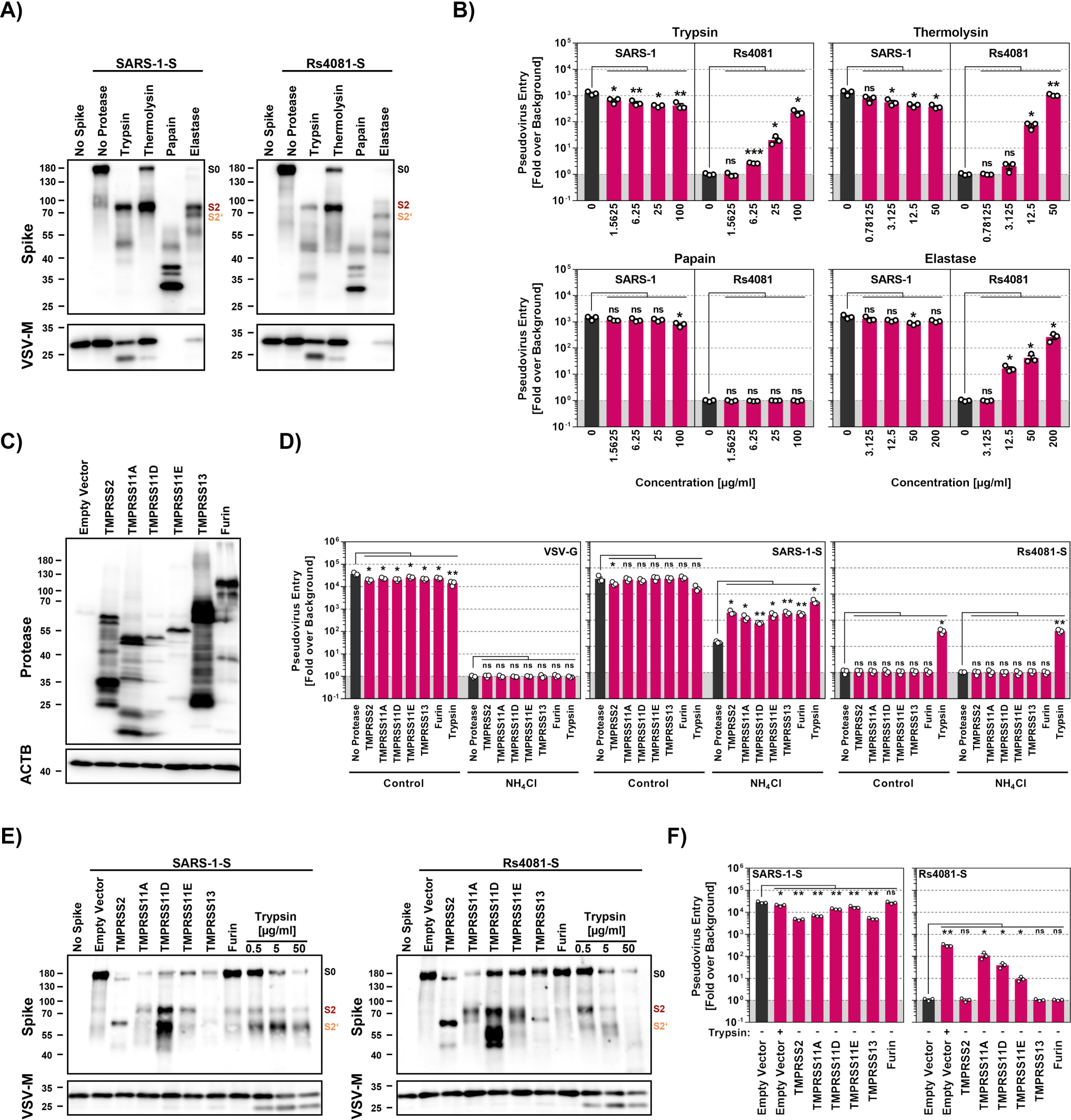
Elastase and type II transmembrane serine proteases can activate the otherwise trypsin-dependent Rs4081 S protein. **A**) Analysis S protein cleavage. Particles pseudotyped with SARS-1-S or Rs4081 S protein (or no S protein) were incubated with the indicated proteases (at highest concentration used for panel B, 20 min incubation) and S protein expression analyzed by immunoblot using an antibody directed against the S2 subunit of SARS-2-S. VSV-M served as loading control. Similar results were obtained in two separate experiments. Bands corresponding to uncleaved S proteins (S0), the S2 subunit (S2) and the S2 subunit cleaved at the S2’ site (S2’) are indicated. **B**) Impact of proteases on cell entry. Particles pseudotyped with SARS-1-S or Rs4081 S protein were treated with the indicated concentrations of trypsin, Thermolysin, papain or elastase for 20 min at 37℃ before addition to Vero cells. The efficiency of S protein-driven cell entry was determined by measuring the activity of virus-encoded firefly luciferase in cell lysates at 16-18h post inoculation. Results for S protein bearing particles were normalized against those obtained for particles bearing no S protein (set as 1). The average (mean) data of three biological replicates are presented, each performed with four technical replicates. Error bars indicate SEM. Statistical significance was assessed by ANOVA (p > 0.05, not significant [ns]; p ≤ 0.05, *; p ≤ 0.01, **; p ≤ 0.001, ***). **C**) Expression of type II transmembrane serine proteases (TTSPs). 293T cells were transiently transfected with plasmids encoding the indicated proteases with a c-myc antigenic tag or empty plasmid and cell lysates were harvested at 48 h after transfection. Cell lysates were analyzed by immunoblot for protease expression using c-myc antibody. Detection of ACTB served as loading control. Similar results were obtained in two separate experiments. **D**) Expression of TTSPs on target cells does not allow for entry driven by the trypsin-dependent Rs4081 S protein. 293T cells transiently expressing the indicated TTSPs of furin were Mock treated or treated with ammonium chloride to block cathepsin L-dependent endo/lysosomal entry and inoculated with pseudotypes bearing SARS-1-S, Rs4081-S or VSV-G. Alternatively, particles were treated with trypsin (50 µg/ml for 30 min) and added to mock treated cells. S-protein-driven cell entry was analyzed by and data presented as described for panel B. The average (mean) data of three biological replicates are presented, each performed with four technical replicates. Error bars show the SEM. Statistical significance was assessed by ANOVA (p > 0.05, not significant [ns]; p ≤ 0.05, *; p ≤ 0.01, **; p ≤ 0.001, ***). **E**) S protein cleavage by TTSPs. Particles pseudotyped with SARS-1-S or Rs4081 S proteins (or no S protein) were produced in 293T cells coexpressing the indicated TTSPs or furin. Alternatively, particles were treated with the indicated concentrations of trypsin for 30 min. S protein expression was analyzed by immunoblot using an antibody directed against the S2 subunit of SARS-2-S. VSV-M served as loading control. Similar results were obtained in two separate experiments. Bands corresponding to uncleaved S proteins (S0), the S2 subunit (S2) and the S2 subunit cleaved at the S2’ site (S2’) are indicated. **F**) Coexpression of TTSPs in particle producing cells can activate the Rs4081 S protein. Particles bearing SARS-1-S or Rs4081 S protein and produced in 293T cells expressing the indicated TTSPs or furin were added to Vero cells. S-protein-driven cell entry was analyzed by and data presented as described for panel B. The average (mean) data of three biological replicates are presented, each performed with four technical replicates. Error bars show the SEM. Statistical significance was assessed by ANOVA (p > 0.05, not significant [ns]; p ≤ 0.05, *; p ≤ 0.01, **; p ≤ 0.001, ***).

Analyses of S protein pseudotyped particles revealed that none of the proteases tested augmented entry driven by SARS-1-S protein and trypsin and thermolysin treatment even reduced particle infectivity (Figure 5B). In contrast, trypsin, thermolysin and elastase allowed for cell entry driven by the Rs4081 S protein in a concentration-dependent manner while papain had no effect (Figure 5B). In sum, Rs4081 S protein can employ elastase, which is expressed in the lung by neutrophils and alveolar macrophages, instead of trypsin for entry into human cells.

### TMPRSS11A, TMPRSS11D and TMPRSS11E cleave coexpressed Rs4081 S protein at the S1/S2 site and confer infectivity to Rs4081 S protein bearing particles

TMPRSS2 and other TTSPs are expressed in the lung and/or gastrointestinal tract and cleave and activate diverse coronavirus S proteins ^12,44,45^. Therefore, we examined whether directed expression of TMPRSS2, TMPRSS11A, TMPRSS11D, TMPRSS11E or TMPRSS13 results in S protein cleavage and promotes entry driven by the Rs4081 S protein. In addition, we analyzed the effect of the expression of furin, which cleaves SARS-2-S at the S1/S2 site in the constitutive secretory pathway of infected cells ^14^.

All proteases examined were efficiently expressed in transfected 293T cells (Figure 5C) and their expression in target cells rescued SARS-1-S but not VSV-G-driven entry from inhibition by ammonium chloride, as expected (Figure 5D). In contrast, protease expression in target cells did not allow for Rs4081 S protein-driven entry (Figure 5D). Therefore, we analyzed whether protease expression in particle-producing cells modulates S protein cleavage and particle infectivity. Expression of TMPRSS11A, TMPRSS11E and furin as well as trypsin-treatment in cells producing SARS-1-S bearing particles had little impact on generation of the S2 fragment (which results from cleavage at the S1/S2 site) (Figure 5E, left panel). Further, TMPRSS11D expression increased production of the S2 fragment and the S2’ fragment (which results from cleavage at the S2’ site) while TMPRSS2 and TMPRSS13 expression and trypsin treatment augmented production of the S2’ fragment and decreased production of the S2 fragment (Figure 5E). Finally, similar findings were made for the Rs4081 S protein, although exposure to 5 and particularly 50 µg/ml trypsin resulted in processing of the S2’ fragment into smaller fragments (Figure 5E).

Expression of TTSPs or furin in particle producing cells or trypsin treatment of particles did not augment cell entry driven by SARS-1-S (Figure 5F). In contrast, expression of TMPRSS11A in particle producing cells increased particle infectivity with similar efficiency as trypsin treatment of particles (Figure 5F). Expression of TMPRSS11D and TMPRSS11E also augmented particle infectivity but with reduced efficiency as compared to TMPRSS11A while expression of TMPRSS2, TMPRSS13 and furin had no effect (Figure 5F). Thus, Rs4081 S protein is cleaved by TMPRSS11A, TMPRSS11D and TMPRSS11E at the S1/S2 site upon protease coexpression and cleavage confers infectivity to Rs4081 S protein-bearing particles.

### Insertion of a multibasic cleavage site increases lung cell infection in a spike-specific fashion

The SARS-CoV-2 S protein but none of the other S proteins studied harbors a multibasic cleavage site at the S1/S2 loop (Figure 6A). The S protein is cleaved at this site by furin and cleavage is essential for robust lung cell entry ^14^. Therefore, we tested whether insertion of the multibasic cleavage site of SARS-2-S in the other S proteins analyzed here increased lung cell entry. The presence of a multibasic cleavage site was compatible with robust expression and particle incorporation of S proteins (Figure 6B) and resulted in efficient proteolytic processing of all S proteins studied (Figure 6B). Notably, the presence of a multibasic cleavage site invariably reduced entry into 293T-ACE2 cells (Figure 6C), which depends on the activity of the S protein activating endo/lysosomal protease cathepsin L. In contrast, the multibasic cleavage site either had no effect or, for the majority of S proteins tested, augmented entry into Calu-3-ACE2 lung cells, with enhancement of entry driven by the S proteins of SARS-CoV-2, RaTG13 and LYRa11 being particularly prominent (Figure 6C). Finally, the presence of a multibasic cleavage site was not sufficient to allow for trypsin-independent 293T-ACE2 or Calu-3-ACE2 cell entry driven by Rs4237 and Rs4081 S proteins (Figure 6C). Thus, a multibasic cleavage site may promote lung cell entry of diverse animal sarbecoviruses but fails to allow for cell entry driven by the S proteins of Rs4237 and Rs4081.

**Fig. 6.**
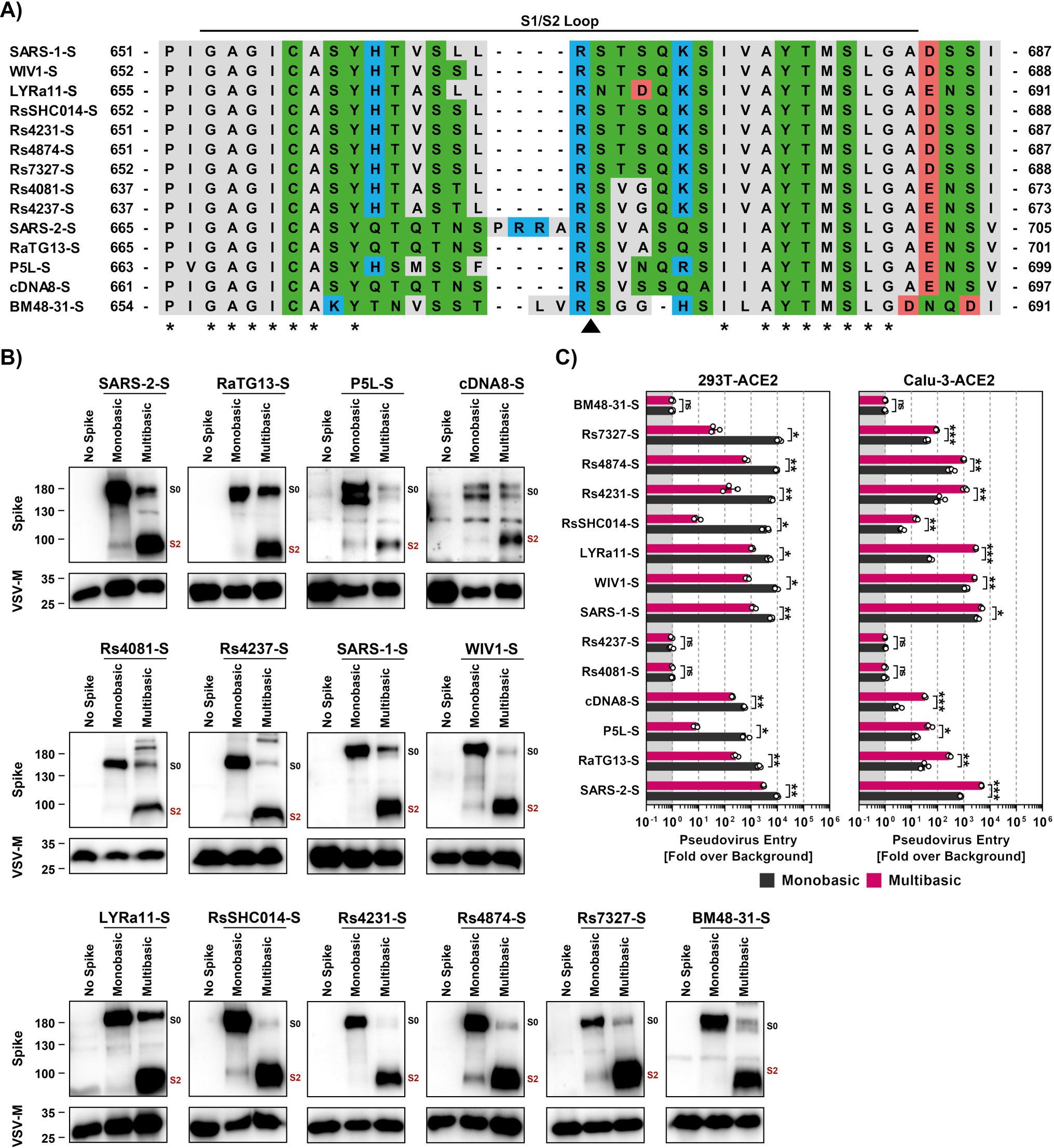
Insertion of a multibasic cleavage site into sarbecovirus S proteins universally increases lung cell entry but does not allow for trypsin-independent entry by RS4081 and Rs4237 S proteins. **A**) Alignment of the S1/S2 loop sequences of the indicated S proteins. Amino acid residues were color coded on the basis of biochemical properties. Asterisks indicate conserved residues. **B**) Analysis of S protein cleavage. Particles pseudotyped with the indicated S proteins were subjected to immunoblot analysis, using anti an antibody directed against the S2 subunit of SARS-2-S. Black and red indicate uncleaved precursor respective S (S0) and S2, respectively. Detection of VSV-M served as a loading control. Shown is a representative immunoblot from three independent experiments. **C**) Impact of the multibasic cleavage site on S protein-driven entry. Particles bearing the indicated S proteins (or no S protein) were added to 293T-ACE2 or Calu-3-ACE2 cells. The efficiency of S protein-driven cell entry was determined by measuring the activity of virus-encoded firefly luciferase in cell lysates at 16-18h post inoculation. Results for S protein bearing particles were normalized against those obtained for particles bearing no S protein (set as 1). Presented are the average (mean) data of three biological replicates, each performed with four technical replicates. Error bars indicate SEM. Statistical significance was assessed by two-tailed Student’s t-tests (p > 0.05, not significant [ns]; p ≤ 0.05, *; p ≤ 0.01, **; p ≤ 0.001, ***).

### The receptor binding domain is a determinant of trypsin-dependent entry of Rs4081

Our studies had so far revealed that Rs4081 and Rs4237 S proteins facilitated entry into human cells only upon pre-cleavage by trypsin or certain other soluble or membrane bound proteases. However, which determinants in the S protein controlled trypsin-dependent entry was unclear. To address this question, we constructed chimeras between SARS-1-S, which facilitates entry in a trypsin-independent fashion, and the Rs4081 S protein, which facilitates entry in a trypsin-dependent fashion. Specifically, we exchanged the S1 subunit between these S proteins or the N-terminal domain (NTD), receptor binding domain (RBD), the domain harboring the S1/S2 and S2’ cleavage sites (priming domain, PD), or NTD jointly with RBD (Figure 7A-B). All chimeric S proteins were efficiently and comparably incorporated into VSV particles (Figure 7C).

**Fig. 7.**
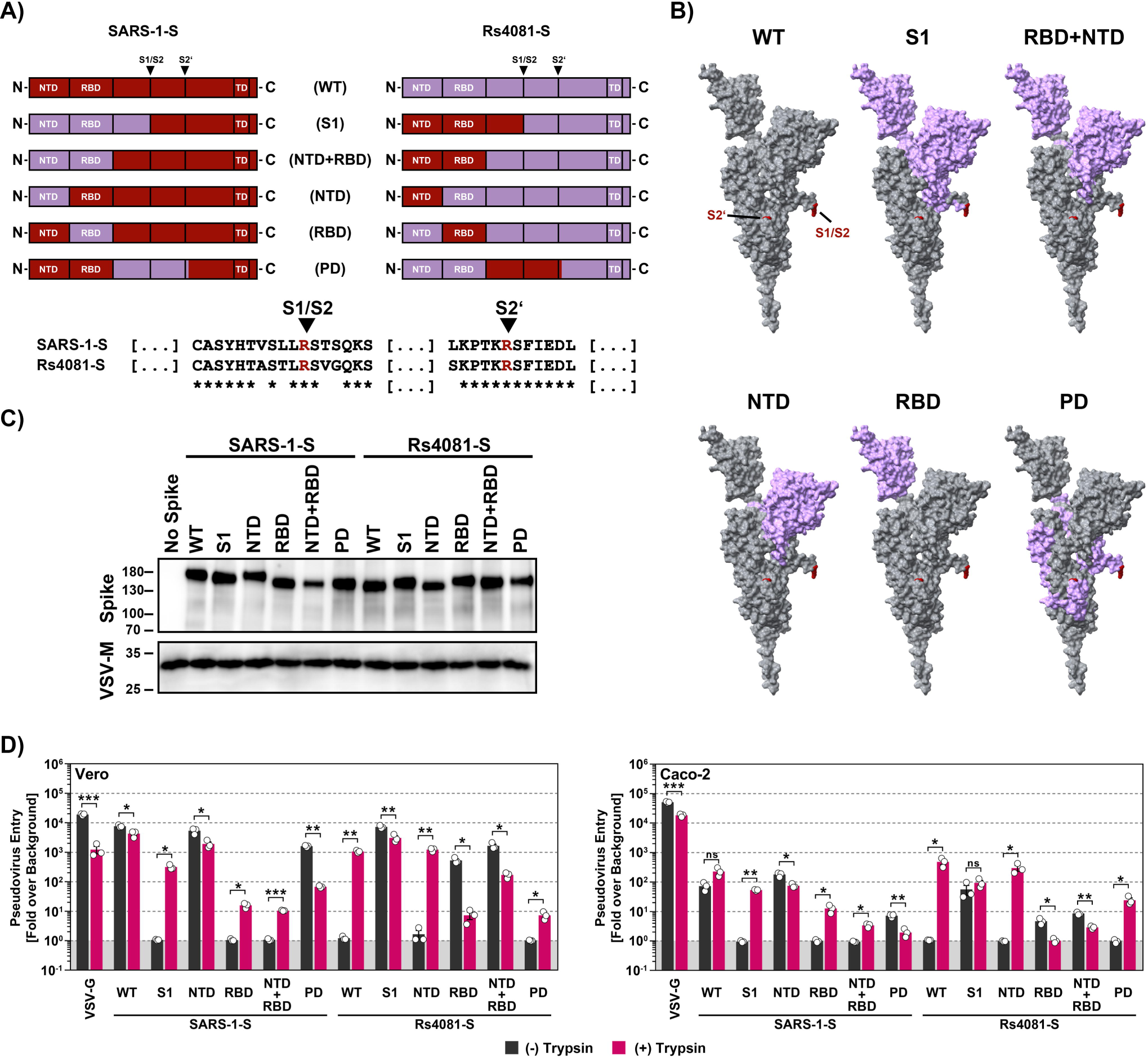
The RBD is the key determinant of trypsin-dependent entry. **A**) Overview of the chimeric SARS-1-S and Rs4081 S proteins analyzed. The sequences of the S1/S2 and S2’ cleavage sites are indicated, asterisk indicate conserved amino acids. **B**) The domains exchanged between SARS-1-S and Rs4081 S proteins are color coded in the context of the S protein monomer. **C**) Expression of chimeric S proteins. Particles pseudotyped with the indicated S protein were subjected to immunoblot analysis, using anti an antibody directed against the S2 subunit of SARS-2-S. Detection of VSV-M served as loading control. Similar results were obtained in two separate experiments. **D**) Cell entry of driven by chimeric S proteins. Particles bearing the indicated S proteins (or no S protein) were treated with trypsin (50 µg/ml for 30 min at 37°C) before addition to Vero or Caco-2 cells. The efficiency of S protein-driven cell entry was determined by measuring the activity of virus-encoded firefly luciferase in cell lysates at 16-18h post inoculation. Results for S protein bearing particles were normalized against those obtained for particles bearing no S protein (set as 1). Presented are the average (mean) data of three biological replicates, each performed with four technical replicates. Error bars indicate SEM. Statistical significance was assessed by two-tailed Student’s t-tests (p > 0.05, not significant [ns]; p ≤ 0.05, *; p ≤ 0.01, **; p ≤ 0.001, ***).

Introduction of the NTD or PD from of Rs4081-S into SARS-1-S was compatible with robust entry into Vero and Caco-2 cells although entry driven by the S protein with PD from the Rs4081 S protein was reduced as compared to WT S protein, and trypsin did not increase entry efficiency (Figure 7D). In contrast, SARS-1-S chimeras harboring the S1 subunit, RBD or NTD+RBD of the Rs4081 S protein mediated entry only upon trypsin treatment. Trypsin-dependent entry mediated by SARS-1-S with the S1 subunit of Rs4081 spike was robust, although not as efficient as entry driven by WT SARS-1-S in the absence of trypsin, while trypsin-dependent entry driven by the SARS-1-S chimera harboring the Rs4081 RBD or NTD+RBD was inefficient (Figure 7D). Finally, the reverse observations were made for Rs4081 S protein harboring domains of SARS-1-S. Entry remained trypsin-dependent when the NTD or PD of SARS-CoV-1 S protein were introduced into Rs4081 S protein while introduction of the S1 subunit, RBD or NTD+RBD allowed for trypsin-independent entry (Figure 7D). In sum, these results show that the RBD is a major determinant of trypsin-dependent entry but also suggest the domains outside the RBD might contribute to this phenotype.

### Trypsin treatment can modulate sarbecovirus neutralization by antibody S2H97

The antibody S2H97 binds to a cryptic epitope within the RBD and recognizes the S proteins of sarbecoviruses from all clades ^46^. The antibody neutralizes particles bearing the S proteins from diverse sarbecoviruses in cell culture and efficiently suppresses SARS-CoV-2 amplification in the lung of experimentally infected hamsters ^46^. Thus, S2H97 and related antibodies could be useful for pandemic preparedness. We investigated whether S2H97 neutralizes particles bearing the S proteins analyzed here and determined whether trypsin treatment modulates neutralization sensitivity. We found that the highest concentration of S2H97 neutralized particles bearing 6 out of the 13 S proteins tested by at least 70% while particles bearing 4 other S proteins were not neutralized (Figure 8). In contrast, entry of particles harboring 2 out of these 4 S proteins was augmented by S2H97 in the absence of trypsin (LYRa11, Rs7327) while moderate but concentration-dependent neutralization was measured in the presence of trypsin (Figure 8).

**Fig. 8.**
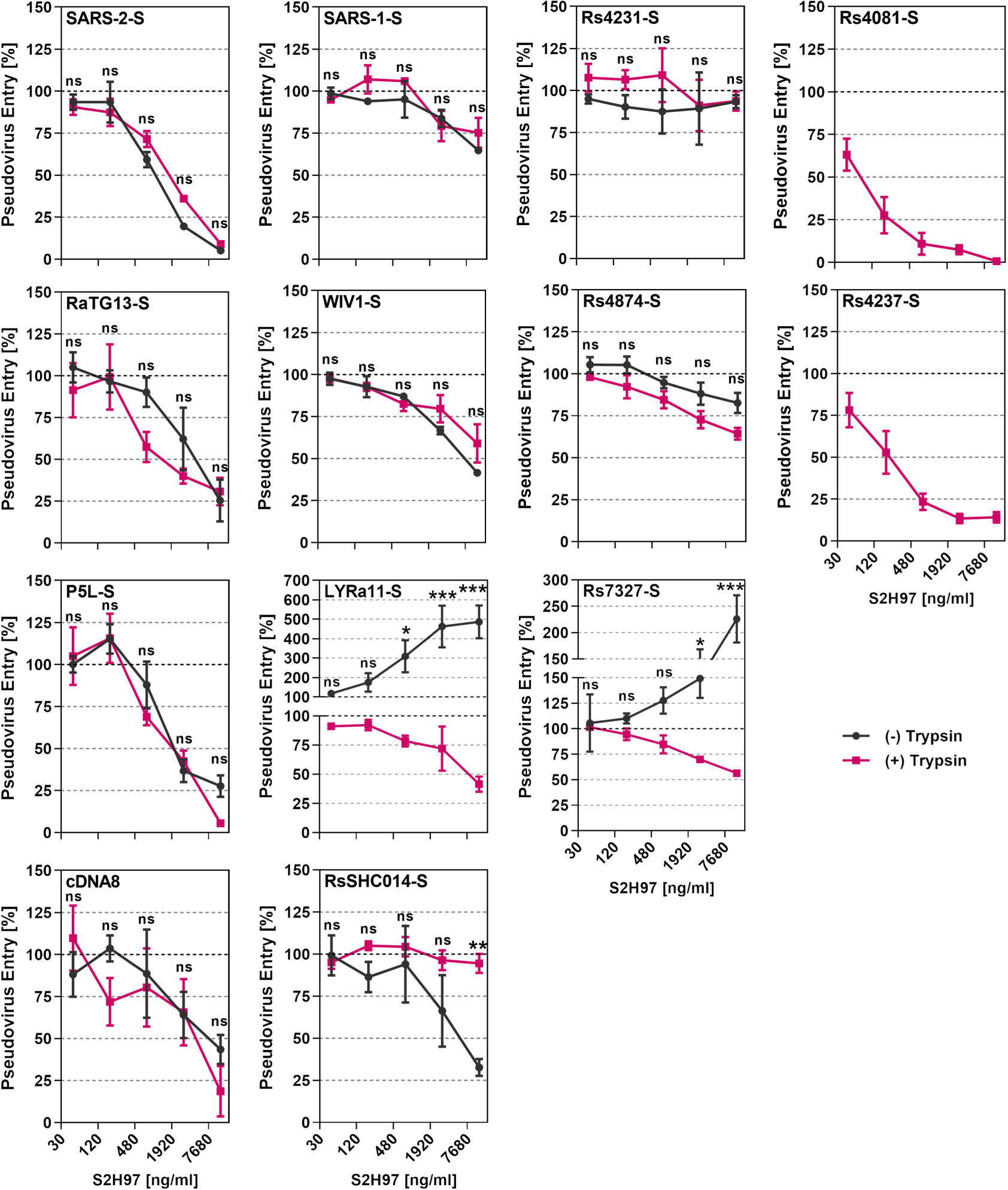
Trypsin treatment modulates sarbecovirus neutralization by the pan-sarbecovirus monoclonal antibody S2H97. Particles bearing the indicated S proteins were preincubated with different concentrations of the pan-sarbecovirus monoclonal antibody S2H97, before being added to Vero-ACE2-TMPRSS2 cells. S protein-driven cell entry was analyzed by measuring the activity of virus-encoded firefly luciferase in cell lysates at 16-18h post inoculation and normalized to entry of in the absence of antibody. Presented are the combined data for 10 plasma. Please see supplementary table 1 for detailed information on the plasma samples. Presented are the average (mean) data of three biological replicates, each performed with four technical replicates. Error bars indicate SEM. Statistical significance was assessed by two-way analysis of variance with Sidak’s multiple comparisons test (p > 0.05, not significant [ns]; p ≤ 0.05, *; p ≤ 0.01, **; p ≤ 0.001, ***).

Finally, trypsin treatment protected particles bearing the RsSHC014 S protein from neutralization by S2H97. These results suggest that S2H97, and potentially related RBD antibodies, neutralize several sarbecoviruses but augment cell entry of others, suggesting limited suitability for pandemic preparedness. Furthermore, our findings indicate that trypsin treatment can alter susceptibility of sarbecoviruses to antibody-mediated neutralization.

### Evidence that antibodies induced upon quadruple vaccination with COVID-19 vaccines cross-neutralize multiple animal sarbecoviruses and that trypsin treatment promotes antibody evasion

The SARS and COVID-19 pandemics demonstrated the massive threat that animal sarbecoviruses pose to human health. However, only few studies systematically analyzed whether immune responses induced by current COVID-19 vaccines may protect against animal sarbecoviruses. Therefore, we determined whether antibodies induced upon infection or vaccination with COVID-19 mRNA vaccines inhibited entry driven by the S proteins analyzed and whether trypsin modulated sensitivity to antibody-mediated neutralization.

Antibodies present in convalescent individuals that were infected by SARS-CoV-2 in the first year of the pandemic efficiently neutralized particles bearing the S proteins of SARS-CoV-2 and the related clade 3 bat sarbecovirus RaTG13, as expected. Robust neutralization was also observed for particles bearing the S proteins of cDNA8, LYRa11, RsSHC014 and Rs4231 while neutralization of particles bearing other S proteins was inefficient (Figure 9A). Similar results were obtained for antibodies induced upon double vaccination (Figure 9A). Finally, particles bearing the S proteins of SARS-CoV-1, WIV1 and Rs4874 were not efficiently neutralized by antibodies induced upon infection or double vaccination but were robustly neutralized by antibodies induced by triple vaccination and, particularly, quadruple vaccination (Figure 9A). In fact, particles bearing all S proteins analyzed were at least 50% neutralized by plasma from quadruple donors, who received three doses of first-generation mRNA vaccines developed against the SARS-CoV-2 B.1 lineage and a fourth dose of a bivalent Omicron BA.5-adapted vaccine. These results suggest that repeated COVID-19 vaccination, including booster vaccination with adapted vaccines, may offer at least partial cross-protection against diverse animal sarbecoviruses.

**Fig. 9.**
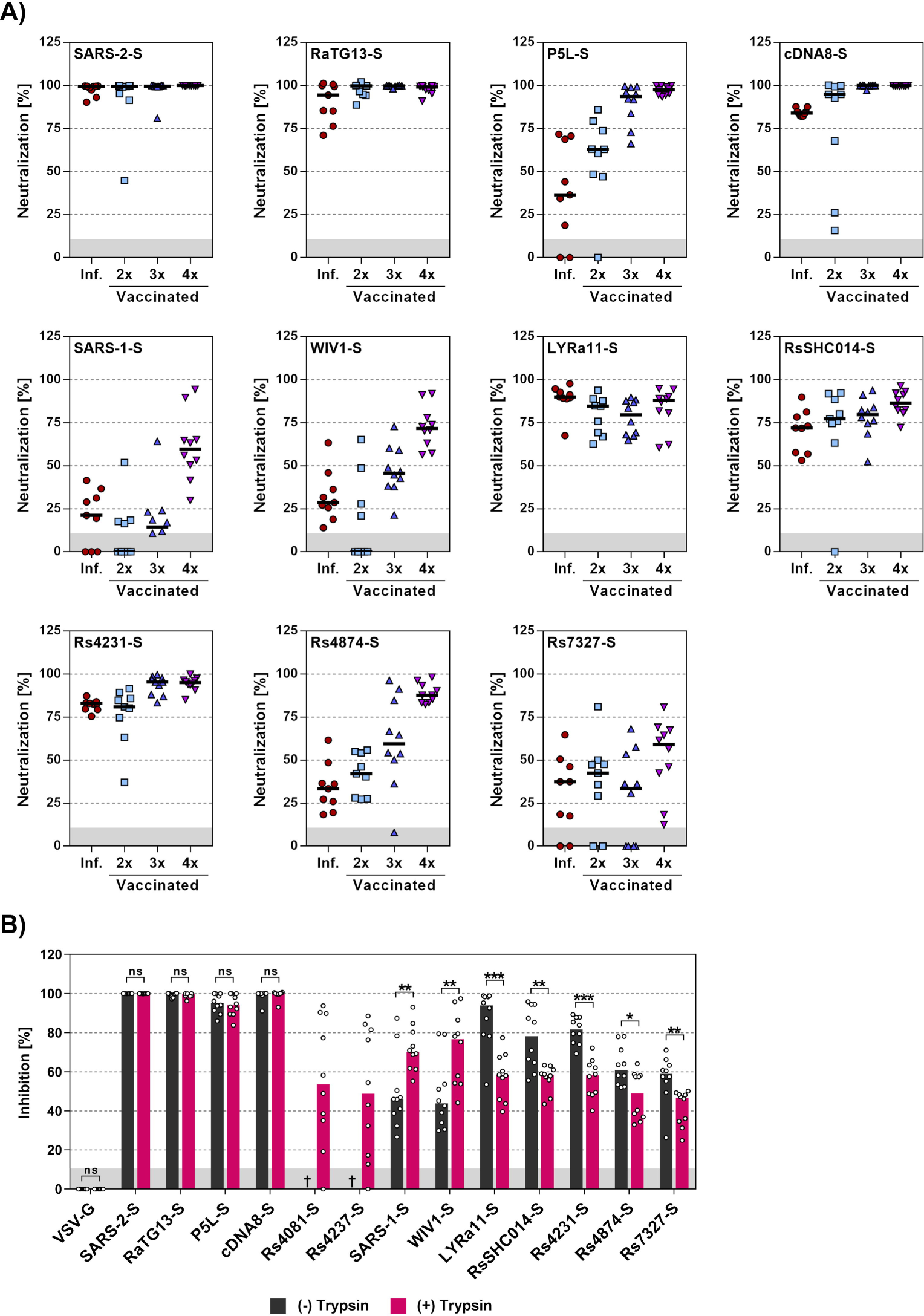
Antibodies induced by quadruple vaccination neutralize particles bearing diverse sarbecovirus S proteins. **A**) Particles bearing the indicated S proteins were preincubated with a 1:25 dilution of plasma from convalescent patients, individuals vaccinated two times with BNT162b2 (BNT/BNT), three times with ChAdOx1-S and BNT162b2 (AZ/BNT/BNT), and four times, including a bivalent, BA.5-adapted booster, before being added to A549-ACE2-TMPRSS2 cells. S protein-driven cell entry was analyzed by measuring the activity of virus-encoded firefly luciferase in cell lysates at 16-18h post inoculation and normalized to entry of in the absence of serum. Presented are the combined data for 10 plasma. Please see supplementary table 1 for detailed information on the plasma samples. **B**) Particles bearing the indicated S proteins were preincubated with trypsin or Mock treated for 30 min before incubation with a fixed 1:25 dilution of serum from triple vaccinated donors that were boostered with a BA.5-adpated vaccine. Subsequently, the particles were added to Vero-ACE2-TMPRSS2 cells. S protein-driven cell entry was analyzed by measuring the activity of virus-encoded firefly luciferase in cell lysates at 16-18h post inoculation and normalized to entry of in the absence of serum. Presented are the combined data for 10 plasma. Please see supplementary for detailed information on the plasma samples tested.

Within the present study, we had found that trypsin promoted entry driven by some S proteins (LYRa11, RsSHC014, Rs7327) and was even essential for entry driven by others (Rs4081, Rs4237) and entry driven by these S proteins was partially (LYRa11, RsSHC014, Rs7327) or fully ACE2-independent (Rs4081, Rs4237) (Figure 3). In contrast, entry driven by a third group of S proteins, comprising SARS-CoV-2, P5L, cDNA8, SARS-CoV-1, WIV1, Rs4231, Rs4874) was not augmented by trypsin (Figure 3) although trypsin rendered entry driven by Rs4874 and Rs4231 S proteins partially ACE2-independent (Figure 3C). We now asked whether differential dependence on trypsin and ACE2 for entry resulted in differential effects of trypsin on antibody-mediated neutralization. Particles bearing S proteins that did not benefit from trypsin for entry and that mediated exclusively ACE2-dependent entry were comparably neutralized in the presence and absence of trypsin (SARS-CoV-2, P5L, cDNA8, Rs4231, Rs4874) or even showed increased neutralization sensitivity in the presence of trypsin (WIV1, SARS-CoV-1) (Figure 9B). In contrast, particles harboring S proteins that facilitated augmented and partially ACE2-independent entry upon trypsin treatment showed reduced neutralization sensitivity in the presence of trypsin (LYRa11, RsSHC014, Rs7327) (Figure 8B). Similarly, particles bearing S proteins Rs4231 and Rs4874 that exhibited no trypsin-dependent augmentation of entry but allowed for partially ACE2-independent entry in the presence of trypsin also showed reduced neutralization upon trypsin treatment. Finally, particles bearing Rs4081 or Rs4237 S proteins that depend on trypsin for entry were 50% neutralized (Figure 9B). In sum, these results suggest that trypsin-dependent usage of an ACE2-independent entry pathway may result in slightly reduced susceptibility to neutralization by antibodies induced upon infection or vaccination.

## Discussion

The spillover of coronaviruses from bats to humans is responsible for the severe respiratory diseases SARS, MERS and likely COVID-19, which emerged within the last two decades ^13,47–49^. Further, two globally circulating endemic human coronaviruses, human coronavirus (HCoV) NL63 and HCoV-229E, which cause the common cold, are believed to have originated from bats and to have caused pandemics in the past ^50^. Therefore, identifying determinants that govern whether animal sarbecoviruses can jump species barriers is an important task.

Our study shows that raccoon dog ACE2 exerts broad receptor activity for animal sarbecoviruses and that ACE2-dependent entry into human lung cells is augmented by the insertion of a multibasic cleavage site into the S protein. Further, we demonstrate that trypsin treatment can confer infectivity of certain animal sarbecoviruses for human cells. Entry of these viruses is ACE2-independent and trypsin-dependent and the latter phenotype is determined by the RBD, confirming previous studies conducted with smaller numbers of spike proteins ^21,31–33^.

Further, we found that TMPRSS2-related proteases, in particular TMPRSS11A and TMPRSS11D, which are known to be expressed in respiratory epithelium, activate ACE2-independent spike proteins for trypsin-independent entry and might thus promote viral invasion of the respiratory tract. Finally, antibodies induced upon quadruple COVID-19 vaccination robustly neutralized entry driven by all S proteins studied and might thus install appreciable protection against zoonotic animal sarbecoviruses. However, usage of the ACE2-independent, trypsin-dependent pathway significantly reduced neutralization sensitivity, which is noteworthy considering that a subset of ACE2-dependent viruses could switch to the ACE2-independent entry pathway in the presence of trypsin.

Our observation that several sarbecovirus S proteins can use ACE2 for entry into human cells is in keeping with previous studies ^19,20,23–27,51,52^ and with the concept that ACE2-binding RBDs evolved independently at least three times, resulting in the SARS-CoV-1 and SARS-CoV-2 clades of Asian origin and the clade comprising SARS-like (SL)-CoVs of European and African descent ^53^. Moreover, a recent study shows that even a MERS-CoV-related bat virus uses ACE2 for entry ^54^ and genetic analysis revealed that the ACE2 gene is under positive selection in bats and primates and might be shaped by pandemic coronaviruses ^55^. Our finding that among animal ACE2 orthologues raccoon dog ACE2 was most efficient at mediating cell entry driven by diverse sarbecovirus S proteins is in keeping with similar findings made for SARS-CoV-1 ^56^. Moreover, this finding highlights that raccoon dogs, which served as intermediate host for SARS-CoV-1 ^57^, might be susceptible to infection by diverse animal sarbecoviruses, although it should be stated that several post entry barriers to sarbecovirus infection have been described ^58^. Indeed, raccoon dogs were found to be susceptible to experimental SARS-CoV-2 infection and transmission of SARS-CoV-2 from experimentally infected to uninfected animals has been observed ^59^. Further, other coronaviruses were recently detected in raccoon dogs that might present a zoonotic threat ^60^. Finally, raccoon dog DNA was associated with SARS-CoV-2 RNA in cages at the Huanan Seafood Wholesale market (DOI 10.5281/zenodo.7754298), the proposed early epicenter of the COVID-19 pandemic ^11^, suggesting that raccoon dogs might have also served as intermediate host for SARS-CoV-2 transmission from reservoir animals to humans.

Results obtained initially in the context of SARS-CoV-1 research revealed that certain sarbecovirus S proteins fail to mediate entry into cells, indicating that they may be non-functional. However, studies in the recent years changed this perception by demonstrating that trypsin treatment can allow certain sarbecovirus S proteins to mediate entry into cell lines that are otherwise refractory ^21,31–33^ and similar findings were reported for other coronavirus S proteins ^35^.

The present study confirms and extends these findings. Thus, using Rs4081 S protein as model, we show that trypsin acts on viral particles but not target cells to facilitate entry into otherwise refractory cell lines and that trypsin cleaves diverse S proteins (including Rs4081-S), producing the S2 and the S2’ fragment, which is associated with membrane fusion. Furthermore, we demonstrate that elastase, a secreted protease that plays a role in several lung diseases ^42,43^, can cleave SARS-1-S and Rs4081 S protein and allow for trypsin-independent entry of Rs4081 S protein-bearing particles, and that the same is true for the respiratory tract expressed TTSPs TMPRSS11A, TMPRSS11D and TMPRSS11E upon expression in particle producing cells. In contrast, expression of these proteases in target cells did not allow for entry. Similarly, TMPRSS2, which is employed by SARS-2-S for lung cell entry ^12,61^, failed to functionally replace trypsin in the context of Rs4081 S protein-driven entry, irrespective of its expression in particle-producing or target cells, and the latter is in keeping with published data ^21^. Collectively, our results are in keeping with the concept that trypsin might promote bat sarbecovirus spread in the perceived central target organ of the bat host, the gastrointestinal tract, but also indicates that several membrane-associated or secreted proteases might allow for infection of the human respiratory tract, potentially promoting zoonotic spillover.

Insertion of a furin cleavage site promoted Calu-3 lung cell entry of ACE2-dependent S proteins, highlighting that optimization of the S1/S2 site may increase human lung cell infection and thus the zoonotic potential of animal sarbecoviruses. However, insertion of a furin cleavage site into the Rs4081 S protein was insufficient for trypsin-independent entry, highlighting that acquisition of a multibasic cleavage site does not universally increase zoonotic potential of animal sarbecoviruses. Instead, mutagenic analysis revealed that the RBD was the key determinant of trypsin-dependent entry, in agreement with previous studies ^21,31^, and it will be interesting to determine whether the RBD is cleaved and whether cleavage is required for trypsin-dependent entry. In this context, it is noteworthy that a previous study indicated that trypsin-treatment decreased rather than increased RBD binding to cells ^21^. Finally, the cellular receptor(s) allowing for trypsin-dependent entry into cells remain(s) to be identified, with known coronavirus receptors playing no role in this process ^21^. Our finding that particles bearing Rs4081 or Rs4237 S protein, which facilitated cell entry only in the presence of trypsin, exhibited marked differences in cell line tropism indicates that more than one receptor might be involved.

Neutralizing monoclonal antibodies could help to contain zoonotic transmission and subsequent human-human spread of animal sarbecoviruses. The antibody S2H97 was found to bind S proteins from sarbecoviruses from all clades and to exert neutralizing activity in cell culture and protect hamsters from viral challenge ^46^. Our analyses confirm that S2H97 neutralizes diverse sarbecoviruses although the inefficient neutralization of particles bearing SARS-CoV-1 S protein was unexpected ^46^. However, S2H97 augmented entry driven by the S proteins of LYRa11 and Rs7327 and antibody-dependent enhancement may increase viral spread and pathogenesis. The underlying mechanism remains to be elucidated but these findings indicate that S2H97 and potentially related antibodies are of limited use for pandemic preparedness. Finally, trypsin treatment sensitized particles bearing the S proteins of LYRa11 and Rs7327 to neutralization by S2H97 while the reverse effect was observed for particles bearing RsSHC014 S protein, suggesting that the availability of the ACE2-independent, trypsin-dependent entry pathway can be modulated neutralization by monoclonal antibodies.

It has previously been appreciated that COVID-19 vaccines can induce antibodies that at least partially neutralize selected animal sarbecoviruses ^27,62–66^. However, systematic analyses are lacking. The present study demonstrates that quadruple vaccination including a bivalent Omicron BA.5-adapted booster induced antibodies that appreciably cross-neutralized particles bearing all S proteins tested. Thus, repeated vaccination might not only come at the benefit of efficient protection against severe COVID-19 but might also provide substantial protection against diverse animal sarbecoviruses. Notably, we obtained evidence that switching to the trypsin-dependent, ACE2-independent entry route reduces neutralization sensitivity, in agreement with the finding that ACE2-independent entry of SARS-CoV-2 conferred by mutation E484D allowed for resistance against a neutralizing antibody ^67^. Collectively, we identified viral and cellular determinants required for animal sarbecovirus infection of human lung cells that may help to predict and combat future spillover events.

## Methods

### Cell culture

All cell lines were incubated in a humidified atmosphere at 37 °C containing 5% CO_2_. 293T (human, kidney; ACC-635, DSMZ), Huh-7 (human, liver; JCRB0403, JCRB; kindly provided by Thomas Pietschmann, TWINCORE, Centre for Experimental and Clinical Infection Research, Hannover, Germany), NCI-H522 (human lung; CRL-5810, ATCC; RRID: CVCL_1567), Vero (African green monkey, kidney; CRL-1586, ATCC; kindly provided by Andrea Maisner, Institute of Virology, Philipps University Marburg, Marburg, Germany), BHK-21 (Syrian hamster, kidney; Laboratory of Georg Herrler, CCL-10, ATCC; RRID: CVCL_1915), PipNi/3 (Common pipistrelle, kidney; RRID: CVCL_RX21), and MyDauLu/47 cells (Daubenton’s bat, lung; RRID: CVCL_RX49) were incubated in Dulbecco’s modified Eagle medium (DMEM, PAN-Biotech) supplemented with 10% fetal bovine serum (FCS, Biochrom), 100 U/ml of penicillin and 0.1 mg/ml of streptomycin (PAN-Biotech). Caco-2 (human, intestine; HTB-37, ATCC, RRID: CVCL_0025) were cultivated in minimum essential medium (GIBCO) supplemented with 10% FCS, 100 U/ml of penicillin and 0.1 mg/ml of streptomycin (PAN-Biotech), 1x non-essential amino acid solution (from 100x stock, PAA) and 10 mM sodium pyruvate (Thermo Fisher Scientific). Calu-3 cells (human, lung; HTB-55, ATCC; kindly provided by Stephan Ludwig, Institute of Virology, University of Münster, Germany) were cultivated in DMEM/F-12 medium (GIBCO) supplemented with 10% FCS, 100 U/ml of penicillin and 0.1 mg/ml of streptomycin (PAN-Biotech), 1x non-essential amino acid solution and 10 mM sodium pyruvate. A549 cells (human, lung; CRM-CCL-185, ATCC), 293T and Calu-3 cells were transduced with murine leukemia virus-based transduction vectors encoding ACE2 and subsequently selected with puromycin (Invivogen), resulting in cell lines that stably expressed ACE2. A549-ACE2 cells were further transduced with murine leukemia virus-based transduction vectors encoding TMPRSS2 and subsequently selected with blasticidin (Invivogen) to obtain A549-ACE2+TMPRSS2 cells. Vero cells stably expressing TMPRSS2 were generated by retroviral transduction and blasticidin-based selection. Vero-TMPRSS2 cells were further transduced with murine leukemia virus-based transduction vectors encoding ACE2 and subsequently selected with puromycin (Invivogen) to obtain Vero-ACE2+TMPRSS2 cells. Cell lines were authenticated using various methods, including STR-typing (human cell lines), amplification and sequencing of a fragment of the cytochrome c oxidase gene, microscopic examination and evaluation of their growth characteristics. All cell lines were regularly tested for mycoplasma.

### Plasmids

Expression plasmids for vesicular stomatitis virus glycoprotein (VSV-G), SARS-1-S Frankfurt-1 (GenBank: AY291315; with a C-terminal truncation of 18 amino acid residues), SARS-2-S (codon-optimized, based on the Wuhan/Hu-1/2019 isolate; with a C-terminal truncation of 18 amino acid residues) were previously described ^12,68^. The sequences of the following spike proteins, bat SARSr-CoV Rs4081-CoV-S (GenBank: KY417143.1), bat SARSr-CoV Rs4237-CoV-S (GenBank: KY417147.1), bat SARSr-CoV WIV1-CoV-S (GenBank: KF367457.1), bat SARSr-CoV LYRa11-CoV-S (GenBank: KF569996.1), bat SARSr-CoV RsSHC014-CoV-S (GenBank: KC881005.1), bat SARSr-CoV Rs4231-CoV-S (GenBank: KY417146.1.), bat SARSr-CoV Rs4874-CoV-S (GenBank: KY417150.1), bat SARSr-CoV Rs7327-CoV-S (GenBank: KY417151.1), bat SARSr-CoV BM48-31/BGR/2008 (GenBank: GU190215.1) were obtained from NCBI (National Library of Medicine) database (https://www.ncbi.nlm.nih.gov/). In addition, the following sequences were obtained from Global Initiative on Sharing All Influenza Data (GISAID) database: SARS-like coronaviruses bat SARSr-CoV RaTG13-CoV-S (codon-optimized, based on the EPI_ISL_402131|2013-07-24), pangolin SL-P5L-CoV-S (EPI_ISL_410540|2017), pangolin SL-cDNA8-CoV-S (EPI_ISL_471461|2019). The sequences were synthesized (Sigma-Aldrich) and inserted into the pCG1 expression vector (kindly provided by Roberto Cattaneo, Mayo Clinic College of Medicine, Rochester, MN, USA) using the BamHI and XbaI restriction sites. The sequences encoding the 18 C-terminal amino acids of these S proteins were removed by PCR-based mutagenesis. For generation of sarbecovirus S proteins harboring a multibasic S1/S2 cleavage site, the amino acid forming the respective S1/S2 regions of animal sarbecovirus S proteins were replaced by the corresponding region of SARS-2-S (amino acid residues 667-701) by overlap-extension PCR. In addition, chimeric S proteins harboring different domains of SARS-1-S and Rs4081-S were constructed by overlap-extension PCR. For generation of expression of plasmid for ACE2 orthologues, the coding sequence for red fox ACE2, palm civet ACE2, Malayan pangolin ACE2^69^, human ACE2, horseshoe bat (*Rhinolophus landeri, R.sinicus, R.affinis*) ACE2, cat ACE2, pig ACE2, raccoon dog ACE2, American mink ACE2 containing C-terminal c-myc-epitope tag were introduced into the pQCXIP plasmid ^70^ via the NotI and PacI restriction sites. Furthermore, we produced an expression plasmid encoding a soluble variant of human ACE2, which was fused to the Fc portion of human immunoglobulin G (sol-hACE2-Fc). For this, we employed PCR amplification to derive the sequence encoding the ACE2 ectodomain, encompassing amino acid residues 1-733, which was subsequently inserted into the pCG1-Fc plasmid ^71^ (kindly provided by Georg Herrler, University of Veterinary Medicine, Hannover, Germany) via PacI and SalI restriction sites. The integrity of all PCR-amplified sequences was confirmed through commercial sequencing services (Microsynth Seqlab).

### Phylogenetic analysis

Phylogenetic analysis (neighbor-joining tree, bootstrap method with 5,000 iterations, Poisson substitution model, uniform rates among sites, complete deletion of gaps/missing data) was performed using the MEGA7.0.26 software. Sequence alignments were performed using the Clustal Omega online tool (https://www.ebi.ac.uk/Tools/msa/clustalo/).

### Production of pseudotyped particles

Rhabdoviral particles bearing coronavirus S proteins, VSV-G or no viral protein (negative control) were prepared according to a published protocol ^72^. In brief, a replication-deficient vesicular stomatitis virus vector that lacks the genetic information for VSV-G and instead codes for two reporter proteins, enhanced green fluorescent protein and firefly luciferase (FLuc), VSV∗ΔG-FLuc (kindly provided by Gert Zimmer, Institute of Virology and Immunology, Mittelhäusern, Switzerland) ^73^ used for particle production. Thus, 293T cells transfected with plasmids encoding the desired viral glycoproteins were inoculated with VSV∗ΔG-FLuc for 1 h at 37 °C, the inoculum was removed and cells were washed with PBS. Of note, for experiments addressing the impact of S protein cleavage by TTSPs during pseudovirus production, cell were cotransfected with TMPRSS2, TMPRSS11A, TMPRSS11D, TMPRS11E, TMPRSS13, or Furin. Subsequently, culture medium supplemented with anti-VSV-G antibody (culture supernatant from I1-hybridoma cells; ATCC no. CRL-2700; except for cells expressing VSV-G) was added to the cells in order to neutralize residual viral particles. After incubation for 16-18 h, the cell culture supernatant was harvested, cleared from debris by centrifugation at 2,000 x g for 10 min, aliquoted and stored at −80 °C until further use.

### Transduction of target cells

For transduction experiments, target cells were seeded into 96-well plates prior to inoculation with equal volumes of pseudotyped particles. For selected experiments, target cells were transfected to express different ACE2 orthologues or proteases 24h prior to infection. In order to investigate the impact of trypsin on S protein-driven cell entry, pseudotyped particles were treated with different concentrations of trypsin for 30 min, followed by a 10 min incubation with trypsin inhibitor (same concentration as for trypsin). Alternatively, pseudotyped particles were treated with different concentrations of thermolysin, elastase or papain. To determine whether ACE2 was required for entry, Vero-TMPRSS2 cells were incubated with recombinant anti-ACE2 neutralizing antibody (Sino Biologics, Cat: 10108-MM36) for 30 minutes prior to inoculation with pseudotyped particles. In order to study neutralization sensitivity of animal sarbecovirus S proteins to the pan-sarbecovirus monoclonal antibody S2H97, pseudotyped particles were pre-incubated for 30 min with medium containing different concentrations of S2H97, before being inoculated onto Vero-ACE2-TMPRSS2 cells. To assess the ability of patient plasma to block S protein-driven cell entry, pseudotyped particles were pre-incubated for 30 min with medium containing a fixed dilution of or patient plasma (1:25), before being inoculated onto A549-ACE2-TMPRSS2 or Vero-ACE2-TMPRSS2 cells. Transduction efficiency was evaluated at 16-18 h post transduction by removing the culture supernatant and lysing the cells in PBS containing 0.5% triton X-100 (Carl Roth) for 30 min at room temperature. The cell lysates were then transferred into white 96-well plates and FLuc activity was measured using a commercial luciferase substrate (Beetle-Juice, PJK) and recorded using the Hidex Sense plate luminometer (Hidex).

### Production of sol-ACE2-Fc

293T cells were seeded in 6-well plates and transfected with 8 µg of sol-ACE2-Fc expression plasmid per well. At 10 h post-transfection, the medium was replaced and cells were incubated for an additional 38 h. Subsequently, the culture supernatant was collected and centrifuged at 2,000 x g for 10 min at 4 °C to remove cellular debris. The resulting clarified supernatant was loaded onto Vivaspin protein concentrator columns with a 30 kDa molecular weight cut-off (Sartorius) and centrifuged at 4,000 x g and 4 °C until the sample was concentrated by a factor of 100. The concentrated sol-ACE2-Fc was aliquot and stored at −80 °C.

### ACE2 binding

To assess the binding efficacy of S proteins to ACE2, 293T cells were seeded into 6-well plates and transfected with the respective S protein-encoding plasmid using calcium-phosphate precipitation method. Empty plasmid served as control. At 24 h post-transfection, the culture medium was replaced with fresh medium, and the cells were further incubated. At 48 h post-transfection, the culture medium was removed, and the cells were resuspended in PBS, pelleted by centrifugation, incubated with different concentration of trypsin, and washed with PBS containing 1% bovine serum albumin (PBS-B). The cells were then resuspended in PBS-B containing solACE2-Fc at a 1:100 dilution and rotated at 4 °C for 1 hour. The cells were then pelleted, washed and resuspended in PBS-B containing anti-human AlexaFluor-488-conjugated antibody in a 1:200 dilution, and rotated for an additional hour at 4 °C. Finally, the cells were washed with PBS-B and analyzed by flow cytometry using ID7000 Spectral Cell Analyzer. The data were processed using the ID7000 Spectral Cell analyzer software (version 1.1.8.18211, Sony Biotechnology, San Jose, CA, USA).

### Immunoblot

To analyze S protein cleavage and incorporation into pseudotyped particles, the pseudotyped particles bearing S proteins were added onto a 20% (w/v) sucrose cushion and subjected to high-speed centrifugation at 25.000 x g for 120 min at 4°C. (i) For investigating S protein incorporation into particles, the supernatant was removed after centrifugation and the pellet was mixed with equal volume of 2x SDS-sample buffer (0.03 M Tris-HCl, 10% glycerol, 2% SDS, 0.2% bromophenol blue, 1 mM EDTA). (ii) To assess S protein cleavage by trypsin, the supernatant was removed after centrifugation and the pellet and residual volume were vortexed and divided into 4 different tubes. Different concentration of trypsin (0 μg/ml, 0.5 μg/ml, 5 μg/ml, 50 μg/ml) were added to the tubes, which were then incubated at 37°C for 20 min. After incubation, the 2x SDS-sample buffer was added and heated for 10 min at 96 °C before SDS-polyacrylamide gel electrophoresis and immunoblotting. The nitrocellulose membranes were blocked in a solution of 5% skim milk powder dissolved in PBS-T (PBS containing 0.05% Tween-20) for 1 h at room temperature. The membranes were then incubated overnight at 4°C with the primary antibody, which was diluted in skim milk solution. After washing three times with PBS-T, the membranes were probed with peroxidase-conjugated anti-mouse or anti-rabbit antibody for 1 h at room temperature. The membranes were then washed three times with PBS-T and incubated with an in house-prepared enhanced chemiluminescent solution (1 ml of solution A: 0.1 M Tris-HCl [pH 8.6], 250 µg/ml luminol sodium salt; 100 µl of solution B: 1 mg/ml para-hydroxycoumaric acid dissolved in dimethyl sulfoxide [DMSO]; 1.5 µl of 0.3 % H_2_O_2_ solution) before being imaged using the ChemoCam imager along with the ChemoStar Imager Software version v.0.3.23 (Intas Science Imaging Instruments GmbH). The primary antibody used for detection of S protein expression was rabbit anti-S2 (SARS-CoV-2 (2019-nCoV) Spike S2 antibody (Biozol, Cat: SIN-40590-T62, diluted 1:2,000) while mouse anti-VSV matrix protein (Kerafast, Cat: EB0011, diluted 1:2,500) was used for detection of M protein expression.

Peroxidase-coupled goat anti-mouse antibody (Dianova, Cat: 115-035-003, diluted 1:2,500) and goat anti-rabbit antibody (Dianova, Cat: 111-035-003, diluted 1:2,500) were used as secondary antibodies

### Detection of SARS-CoV-2 IgG

We measured SARS-CoV-2 IgG by quantitative ELISA (anti-SARS-CoV-2 S1 Spike protein domain/receptor binding domain IgG SARS-CoV-2-QuantiVac, EUROIMMUN, Lübeck, Germany) according to the manufacturer’s instructions (dilution up to 1:4,000). We used an AESKU.READER (AESKU.GROUP, Wendelsheim, Germany) and the Gen5 2.01 Software for analysis.

### Patient plasma samples

Before analysis, all plasma samples underwent heat-inactivation at 56°C for 30 min and prescreening for robust neutralization of pseudotyped particles bearing SARS-2-S. Convalescent plasma was obtained from COVID-19 patients treated at the intensive care unit of the University Medicine Göttingen (UMG) under approval given by the ethic committee of the UMG (SeptImmun Study 25/4/19 Ü). Plasma from vaccinated individuals was collected at Hannover Medical School under approval given by the Institutional Review Board of Hannover Medical School (8973_BO_K_2020, amendment Dec 2020). Written informed consent was obtained from each participant prior to the use of any plasma samples for research.

### Data normalization and statistical analysis

Data analysis was performed using ID7000 Spectral Cell analyzer software (version 1.1.8.18211, Sony Biotechnology, San Jose, CA, USA), Microsoft Excel (part of the Microsoft Office software package, version 2019, Microsoft Corporation) and GraphPad Prism 8 version 8.4.3 (GraphPad Software). Only *P* values of 0.05 or lower were considered statistically significant (*P* > 0.05, not significant [ns]; *P* ≤ 0.05, *; *P* ≤ 0.01, **; *P* ≤ 0.001, ***). Specific details on the statistical test and the error bars are indicated in the figure legends.

## Data availability

Datasets generated and/or analyzed during the current study are available in the paper or are appended as supplementary data. Source data are provided in this paper.

## Acknowledgements

We thank Anne Cossmann for data collection and curation. SP acknowledges funding by the EU project UNDINE (grant agreement number 101057100), the COVID-19-Research Network Lower Saxony (COFONI) through funding from the Ministry of Science and Culture of Lower Saxony in Germany (14-76103-184, projects 7FF22, 6FF22, 10FF22) and the German Research Foundation (Deutsche Forschungsgemeinschaft, DFG; PO 716/11-1). G.M.N.B. acknowledges funding by German Center for Infection Research (grant no 80018019238), the European Regional Development Funds (Defeat Corona, ZW7-8515131), and Getting AIR (ZW7-85151373) and the Ministry for Science and Culture of Lower Saxony (Niedersächsisches Ministerium für Wissenschaft und Kultur; 14-76103-184, COFONI Network, project 4LZF23. LZ acknowledges funding by the China Scholarship Council (CSC) (202006270031). Funding sources had no role in the design and execution of the study, the writing of the manuscript and the decision to submit the manuscript for publication. The authors did not receive payment by a pharmaceutical company or other agency to write the publication. The authors were not precluded from accessing data in the study, and they accept responsibility to submit for publication.

## Author contributions

Conceptualization: M.H.; Methodology: S.P., and M.H.; Investigation: L.Z., H.H.C., N.K., L.G., and M.H.; Formal analysis: M.H.; Resources: B.H., A.H., S.R.S, H.-M.J., M.V.S., G.M.N.B., M.A.M., C.D., O.M., M.S.W., and Z.H. Q.; Funding acquisition: C.D., G.M.N.B. and S.P.; Writing – original draft: S.P., L.Z., M.H.; Writing – review & editing: all authors.

## Additional information

See Supplementary Information

