## Supplemental information for "ACE2-independent sarbecovirus cell entry is supported by TMPRSS2-related enzymes and reduces sensitivity to antibody-mediated neutralization"

[1]Infection Biology Unit, German Primate Center, 37077 Göttingen, Germany. [2]Faculty of Biology and Psychology, Georg-August-University Göttingen, 37073 Göttingen, Germany. [3]Platform Infection Models, German Primate Center, 37077 Göttingen, Germany. [4]Junior Research Group Herpesviruses, German Primate Center, 37077 Göttingen, Germany. [5]Division of Molecular Immunology, Department of Internal Medicine 3, Friedrich-Alexander University of Erlangen-Nürnberg, 91054 Erlangen, Germany. [6]Department of Rheumatology and Immunology, Hannover Medical School, 30625 Hannover, Germany. [7]German Centre for Infection Research (DZIF), partner site Hannover-Braunschweig, Hannover, Germany. [8]Institute of Virology, Campus Charité Mitte, Charité - Universitätsmedizin Berlin, corporate member of Freie Universität Berlin and Humboldt-Universität zu Berlin, 10117 Berlin, Germany. [9]Department of Anesthesiology, University of Göttingen Medical Center, Göttingen, Georg-August University of Göttingen, Robert-Koch-Straße 40, 37075 Göttingen, Germany. [10]NHC Key Laboratory of Systems Biology of Pathogens, Institute of Pathogen Biology, Chinese Academy of Medical Sciences and Peking Union Medical College, Beijing, China

* (S.P.), (M.H.)

**Supplemental Table 1**

| **Plasma from convalescent donors** | | | | | | | |
|---|---|---|---|---|---|---|---|
| ID | Sex | Age (years) | Infected | Vaccinated | Vaccine | Time since infection (days) | Anti-S1 IgG (BAU/ml) |
| SI15 | M | 65 | yes | no | n.a. | unknown | unknown |
| SI18 | F | 74 | yes | no | n.a. | unknown | unknown |
| SI20 | M | 61 | yes | no | n.a. | unknown | unknown |
| SI22 | F | 25 | yes | no | n.a. | unknown | unknown |
| SI23 | F | 69 | yes | no | n.a. | unknown | unknown |
| SI24 | M | 61 | yes | no | n.a. | unknown | unknown |
| SI27 | M | 52 | yes | no | n.a. | unknown | unknown |
| SI33 | M | 75 | yes | no | n.a. | unknown | unknown |
| SI51 | M | 71 | yes | no | n.a. | unknown | unknown |
| **Plasma from 2x vaccinated donors** | | | | | | | |
| ID | Sex | Age (years) | Infected | Vaccinated | Vaccine | Time since last vaccination (days) | Anti- S1 IgG (BAU/ml) |
| 4847 | F | 57 | no | yes | BNT/BNT | 26 | 381 |
| 4848 | M | 29 | no | yes | BNT/BNT | 26 | 1221 |
| 4849 | M | 39 | no | yes | BNT/BNT | 24 | 2032 |
| 4863 | F | 58 | no | yes | BNT/BNT | 26 | 618 |
| 4865 | M | 57 | no | yes | BNT/BNT | 25 | 747 |
| 4867 | F | 52 | no | yes | BNT/BNT | 30 | 1529 |
| 4868 | F | 56 | no | yes | BNT/BNT | 30 | 392 |
| 4872 | F | 37 | no | yes | BNT/BNT | 25 | 1164 |
| 4877 | F | 29 | no | yes | BNT/BNT | 28 | 1131 |
| **Plasma from 3x vaccinated donors** | | | | | | | |
| ID | Sex | Age (years) | Infected | Vaccinated | Vaccine | Time since last vaccination (days) | Anti- S1 IgG (BAU/ml) |
| 8624 | F | 31 | no | yes | AZ/BNT/BNT | 123 | 2629 |
| 8631 | F | 61 | no | yes | AZ/BNT/BNT | 123 | 5429 |
| 8632 | F | 46 | no | yes | AZ/BNT/BNT | 123 | 2374 |
| 8639 | F | 50 | no | yes | AZ/BNT/BNT | 110 | 5576 |
| 8645 | F | 57 | no | yes | AZ/BNT/BNT | 134 | 8758 |
| 8649 | | 53 | no | yes | AZ/BNT/BNT | 128 | 7828 |
| 8663 | F | 54 | no | yes | AZ/BNT/BNT | 127 | 2644 |
| 8664 | M | 51 | no | yes | AZ/BNT/BNT | 129 | 4819 |
| 8700 | F | 29 | no | yes | AZ/BNT/BNT | 136 | 3610 |

| | | | | | | | |
|---|---|---|---|---|---|---|---|
| 8701 | M | 33 | no | yes | AZ/BNT/BNT | 134 | 8194 |
| **Plasma from 4x vaccinated donors** | | | | | | | |
| ID | Sex | Age (years) | Infected | Vaccinated | Vaccine | Time since last vaccination (days) | Anti- S1 IgG (BAU/ml) |
| 9474 | F | 54 | no | yes | unknown/XBB | 33 | 2629 |
| 9476 | F | 51 | no | yes | unknown/XBB | 33 | 5429 |
| 9477 | M | 52 | no | yes | unknown/XBB | 33 | 2374 |
| 9479 | M | 62 | no | yes | unknown/XBB | 33 | 5576 |
| 9481 | F | 58 | no | yes | unknown/XBB | 23 | 8758 |
| 9484 | M | 45 | no | yes | unknown/XBB | 33 | 7828 |
| 9488 | M | 54 | no | yes | unknown/XBB | 26 | 2644 |
| 9493 | F | 61 | no | yes | unknown/XBB | 28 | 4819 |
| 9494 | F | 50 | no | yes | unknown/XBB | 33 | 3610 |
| 9496 | M | 40 | no | yes | BNT/BNT/ MOD/XBB | 33 | 8194 |

n.a.: not applicable

BNT: BNT162b2, XBB: BNT162b2 omicron XBB.1.5, AZ: Vaxzevria; MOD: Spikevax

### Supplemental Figure 1

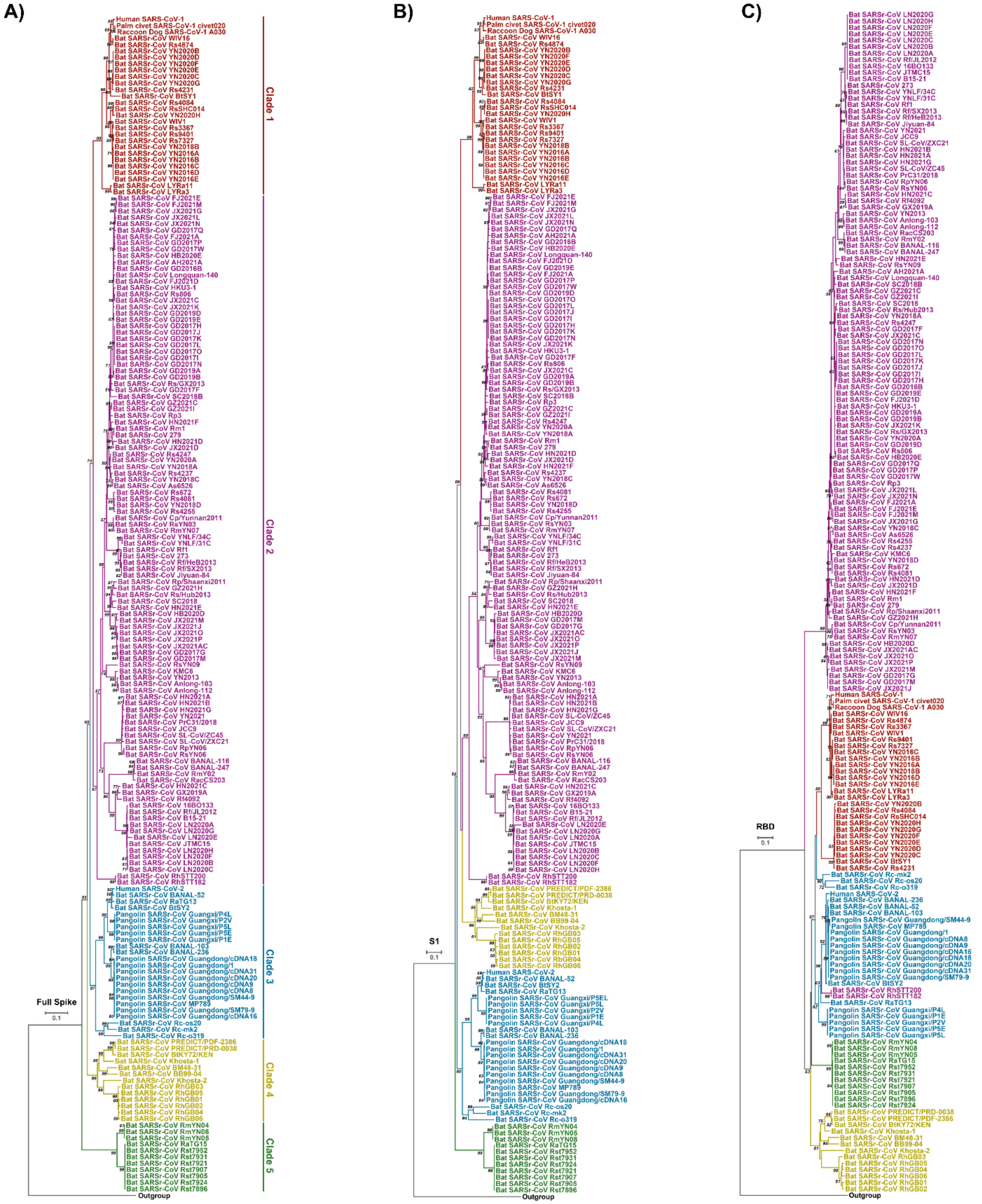

**Supplemental Figure 1. Phylogenetic analysis of sarbecoviruses.**

Phylogenetic analysis was based on full S amino acid sequence (**A**), S1 subunit (**B**) or RBD (**C**).

#### Supplemental Figure 2

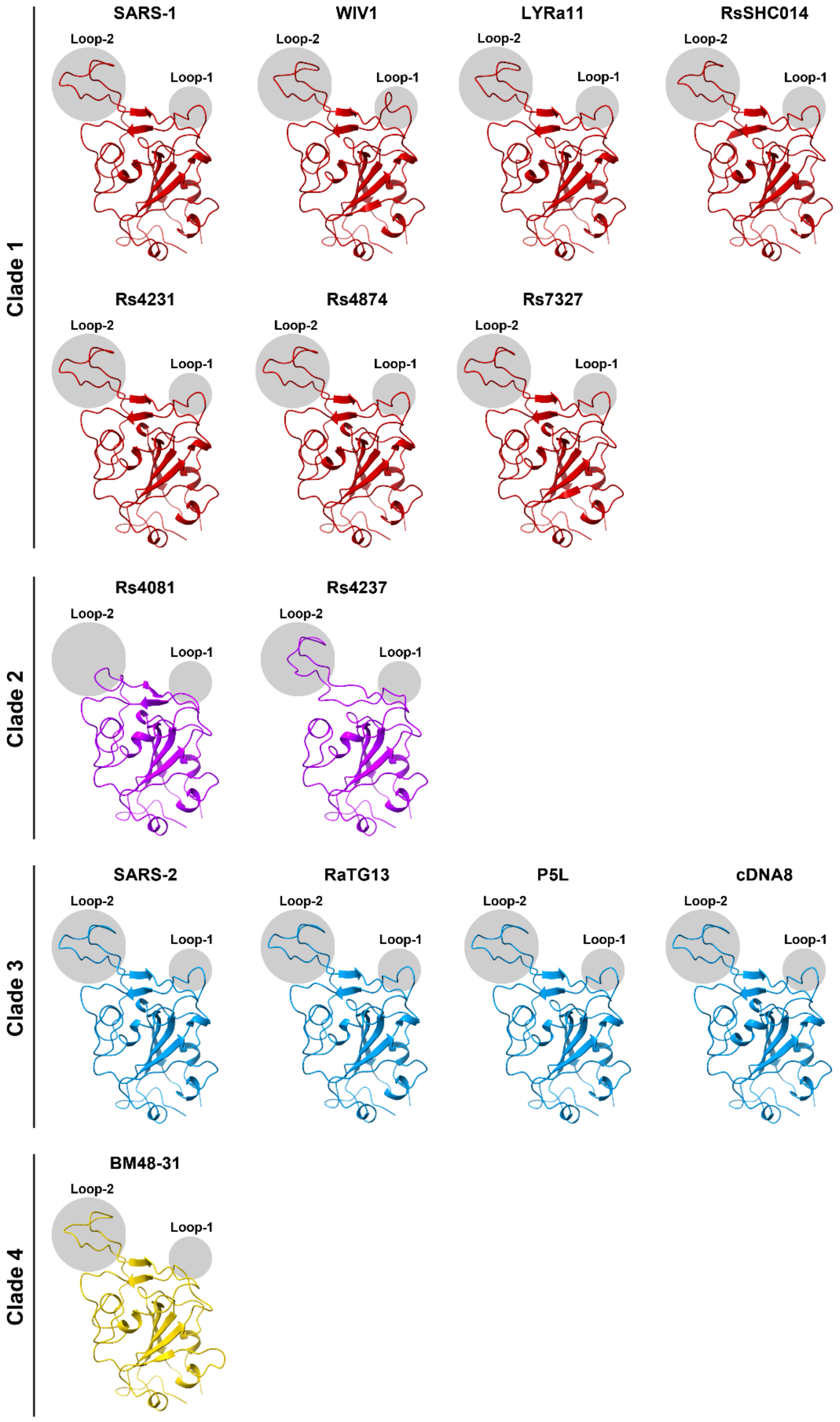

**Supplemental figure S2. Structure of the RBD of sarbecovirus S proteins.**

#### Supplemental Figure 3

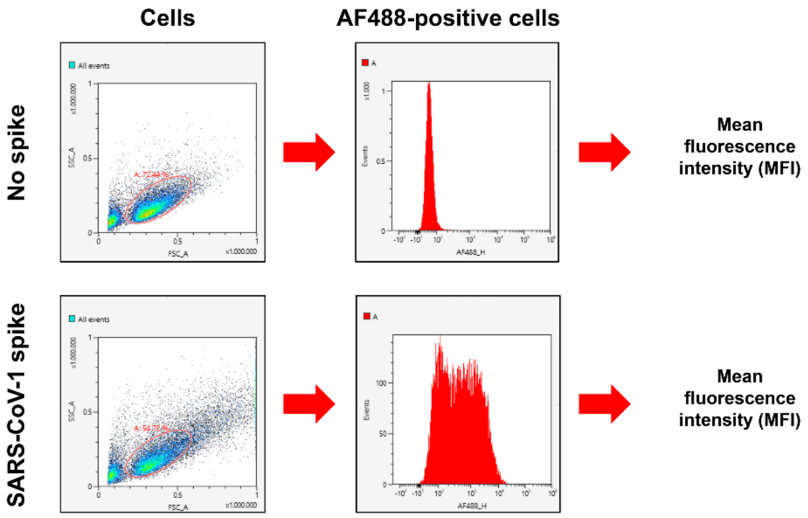

**Supplemental Figure 3. Gating strategy for flow cytometry.**

Gating strategy for flow cytometry (related to Figure 2A).

#### Supplemental Figure 4

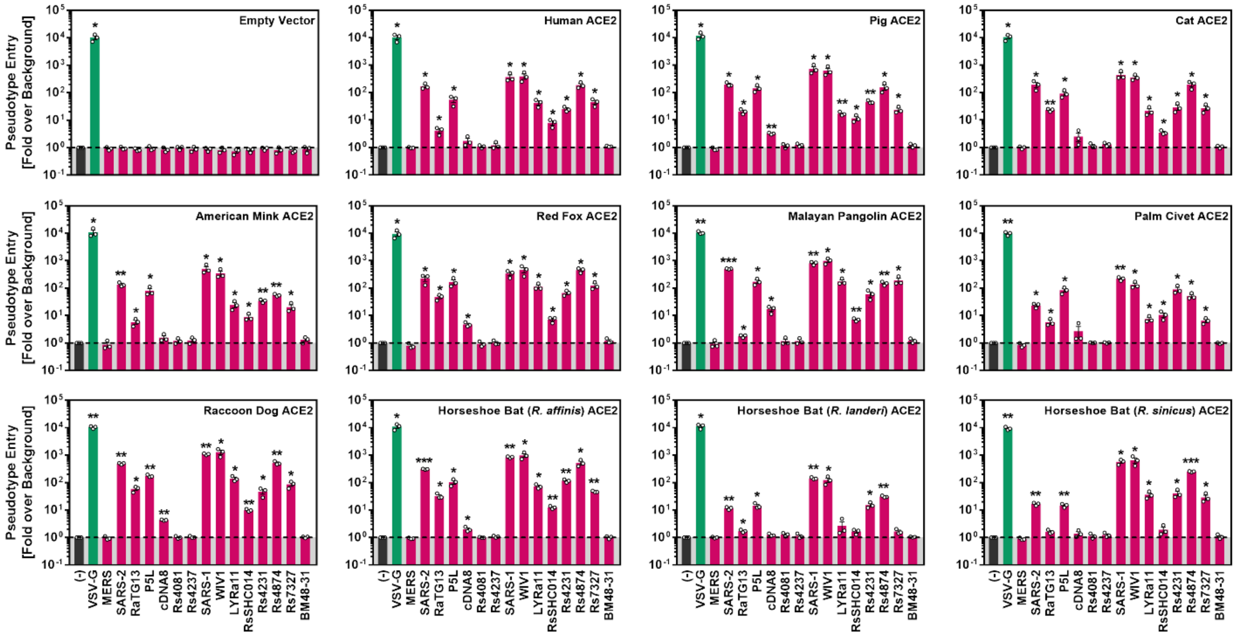

**Supplemental Figure 4. Receptor activity of animal ACE2 orthologues.**

Receptor activity of ACE2 orthologues. BHK-21 cells transiently expressing the indicated ACE2 orthologues (or empty vector) were inoculated with pseudotyped particles bearing the indicated S proteins (or no S protein). Entry into cells expressing ACE2 orthologues was normalized against entry into cells by particles bearing no S protein (set as 1). Particles bearing VSV-G (ACE2-independent entry) served as control. Presented are the average of (mean) data from three biological replicates (each conducted with four technical replicates). Error bars show the SEM. Statistical significance was assessed by two-tailed Student's t-tests ($p > 0.05$, not significant [ns]; $p \leq 0.05$, *; $p \leq 0.01$, **; $p \leq 0.001$, ***).

**Supplemental Figure 5**

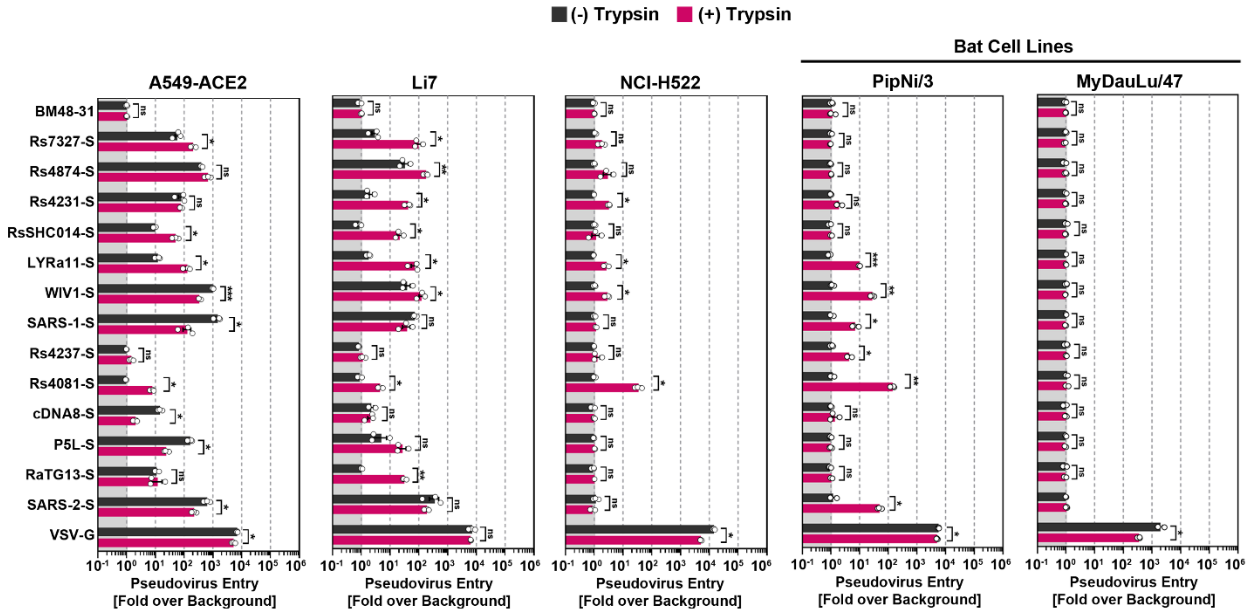

**Supplemental Figure 5. S protein driven cell entry in the presence and absence of trypsin.** Related to figure 3. Particles bearing the indicated S proteins (or no S protein) were preincubated with or without trypsin before being added to the respective cell lines. S protein driven cell entry was analyzed by measuring the activity of virus-encoded firefly luciferase in the cell lysate at 16-18h post inoculation. Presented are the average of (mean) data from three biological replicates (each conducted with four technical replicates) in which cell entry was normalized against that measured for particles bearing no S protein (set as 1). Error bars show the SEM. Statistical significance was assessed by two-tailed Student's t-tests ($p > 0.05$, not significant [ns]; $p \leq 0.05$, *; $p \leq 0.01$, **; $p \leq 0.001$, ***).

**Supplemental Figure 6**

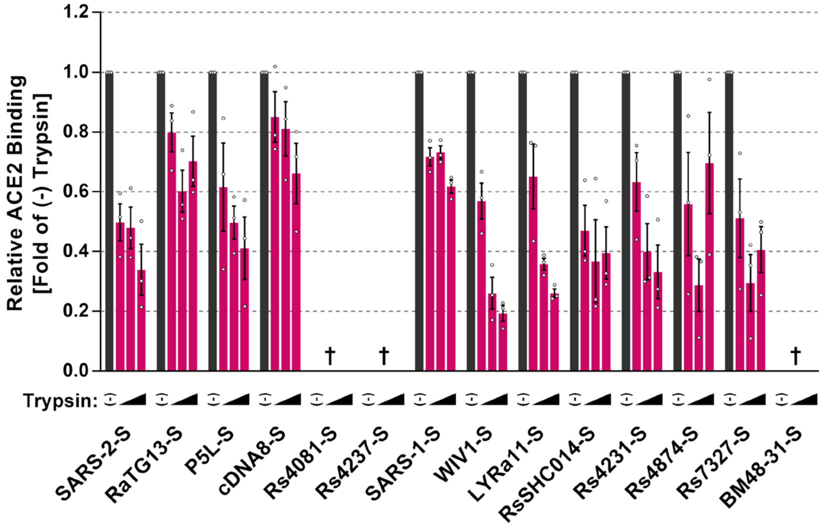

**Supplemental Figure 6. Trypsin does not increase ACE2 binding.**

Binding of soluble human ACE2 to S protein expressing cells. 293T cells transiently expressing the indicated S proteins (or no S protein) were pre-incubated with different concentrations of trypsin or mock incubated. Thereafter, samples were incubated with soluble ACE2 containing a C-terminal Fc-tag (derived from human immunoglobulin G; solACE2-Fc) and subsequently incubated with an AlexaFluor-488-coupled secondary antibody. Finally, solACE2-Fc binding was analyzed by flow cytometry. Presented are the average (mean) data from three biological replicates (each conducted with single samples) in which solACE2-Fc binding to S protein expressing cells was normalized to binding to cells that were not treated with trypsin (set as 1). Error bars indicate SEM.
